# Insights into the role of dopamine in rhizosphere microbiome assembly

**DOI:** 10.1101/2024.08.07.607067

**Authors:** Yezhang Ding, Hunter K. Vogel, Yi Zhai, Hans K. Carlson, Peter F. Andeer, Vlastimil Novak, Nakian Kim, Benjamin P. Bowen, Amber N. Golini, Suzanne M. Kosina, Devin Coleman-Derr, John P. Vogel, Trent R. Northen

## Abstract

Dopamine plays a critical role in animal physiology and interactions with gut microbes. In plants, dopamine is known to function in plant defense and abiotic stress tolerance; however, its role in mediating plant-microbiome interactions remains unexplored. In this study, we observed that dopamine is one of the most abundant exometabolites with natural variation in root exudates across diverse *Brachypodium distachyon* lines, suggesting a potential role in rhizosphere microbial assembly. To further investigate this, we colonized ten natural *B. distachyon* lines with a 16-member bacterial synthetic community (SynCom), collected paired metabolomic and 16S rRNA sequencing data, and performed an association analysis. Our results revealed that dopamine levels in root exudates were significantly associated with the abundance of six SynCom members in a hydroponic system. *In vitro* growth studies demonstrated that dopamine had a significant effect on the growth of the same six bacterial isolates. Additionally, treating soil directly with dopamine enriched Actinobacteria, consistent with both the SynCom-dopamine correlations and the isolate growth results. Collectively, our study underscores the selective influence of dopamine on rhizosphere microbial communities, with implications for precision microbiome management.

## Introduction

In recent years, there has been a growing interest in the field of plant-microbiome interactions, primarily due to their significant potential to enhance agricultural production by fostering plant growth, plant defense, and soil health^1,2^. The rhizosphere is one of Earth’s most complex ecosystems, serving as a hotspot that fosters extraordinary microbial diversity. The composition of microbial communities in the rhizosphere varies with plant developmental stage, plant genotype, and soil environment^3,4,5,6^. Root exudates have been considered key drivers in mediating interactions of plants with rhizosphere microbes^7,8^.

Plant roots exude up to 20% of photosynthetically fixed carbon and 15% of the nitrogen absorbed by plants^8,9^. Root exudates include a diverse array of primary and specialized metabolites as well as complex polymers that shape crucial interactions within the rhizosphere^8,9^. The composition of root exudates is influenced by a combination of plant genetics, soil properties, microbial interactions, and environmental conditions^8,10^. Hydroponic systems are often used for studying root exudate composition and dynamics due to their controlled environment, direct access to exudates, non-destructive sampling methods, precision in experimental setup, and suitability for investigating specific plant-microbe interactions^11,12,13,14^. For example, the exudate dynamics of three phylogenetically distinct plant species, *Arabidopsis thaliana*, *Brachypodium distachyon*, and *Medicago truncatula*, were studied in hydroponics, revealing both significant differences between species and a core metabolome shared by all three species^14^.

Plant specialized metabolites are considered a potential mechanism shaping the plant-microbiome interactions due to their chemical diversity and antimicrobial properties^15,16^. Recent investigations have revealed that plant specialized metabolites, such as coumarin, camalexin, triterpenes, flavonoids, and benzoxazinoids, play a crucial role in mediating interactions between plants and the soil microbiome^16,17^. Pathway mutants are often used to discover the relationship between plant specialized metabolites and the root microbiome^17^. However, the utilization of genetic mutants relies on preexisting knowledge of the biosynthetic pathways and tends to be biased towards the study of readily identifiable candidate metabolites. In contrast, employing natural variation in natural lines allows for the relatively unbiased discovery of unexpected links between root exometabolites and the rhizosphere microbial community.

Catecholamines, such as dopamine and norepinephrine, are known to affect the human gut microbial growth^18,19,20,21^. These metabolites can be produced by commensal gut microorganisms and also derived from dietary sources^22^. Different human bacteria respond differently when exposed to these metabolites^20^. In addition, catecholamines can influence the expression of bacterial genes involved in virulence, stress response, and biofilm formation, thereby impacting the overall growth and function of the gut microbiota^23,24^. Many plant species also produce dopamine and its related compounds^25^. In plants, dopamine can promote plant disease resistance and enhance plant tolerance against abiotic stresses^26^. Recently, dopamine was shown to be present in the spent medium of the model grass *B. distachyon*^27^. However, its role in mediating plant-microbiome interactions has not been explored.

To uncover specialized metabolites involved in plant-microbiome interactions in *B. distachyon*, we inoculated diverse lines with a 16-member bacterial SynCom in a hydroponic system, collected paired metabolomic and 16S rRNA sequencing data, and conducted an association analysis. We found that the level of dopamine, one of the most abundant exometabolites, varied in root exudates of diverse *Brachypodium* lines. Association analysis with paired metabolomics and 16S rRNA sequencing data revealed that dopamine levels in root exudates were significantly associated with the abundance of two Actinobacteria and four Proteobacteria. Consistently, *in vitro* growth studies demonstrated that dopamine had a direct growth effect on the six SynCom bacteria. Furthermore, treating soil directly with dopamine specifically enriched Actinobacteria, supporting both the SynCom-dopamine correlations and the isolate growth results. Taken together, these findings unveil a mechanism through which plants use dopamine to modulate soil microbial communities.

## Results

### Natural variation in root exudates modulates a SynCom composition

To investigate the effects of natural variation in root exudates on plant-microbiome interactions, we selected ten *B. distachyon* lines that are the parental lines for a recombinant inbred line (RIL) population known to exhibit a large degree of phenotypic diversity^28^. These genetically and phenotypically diverse inbred *B. distachyon* lines were grown in a hydroponics-based system (for the workflow see Supplementary Fig. 1), with a 16-member bacterial SynCom isolated from a *Panicum virgatum* (switchgrass) rhizosphere (Supplementary Table 1)^29^. After an additional three weeks of incubation, we observed significant increases in the final OD_600_ of the plant spent medium as compared to that of the control (No Plants) (Supplementary Fig. 2). Consistently, we noted that these hydroponically grown *B. distachyon* lines displayed distinct growth phenotypes both in the absence and presence of the SynCom (Supplementary Fig. 3).

We next examined how host genetic diversity shapes rhizosphere community composition. To do this, we performed 16S rRNA gene profiling on the rhizosphere communities, which revealed differential relative abundances of the SynCom members in the plant spent medium of these *B. distachyon* lines (Fig. 1a, Supplementary Table 2). Notably, one bacterial strain, *Mucilaginibacter* OAE612, was not recovered by 16S rRNA gene amplicon sequencing, possibly due to unsuitable growth conditions for the bacterium. Principal coordinate analysis further revealed distinct microbiome profiles in the plant spent medium of these *B. distachyon* lines (Fig. 1b).

**Fig. 1.**
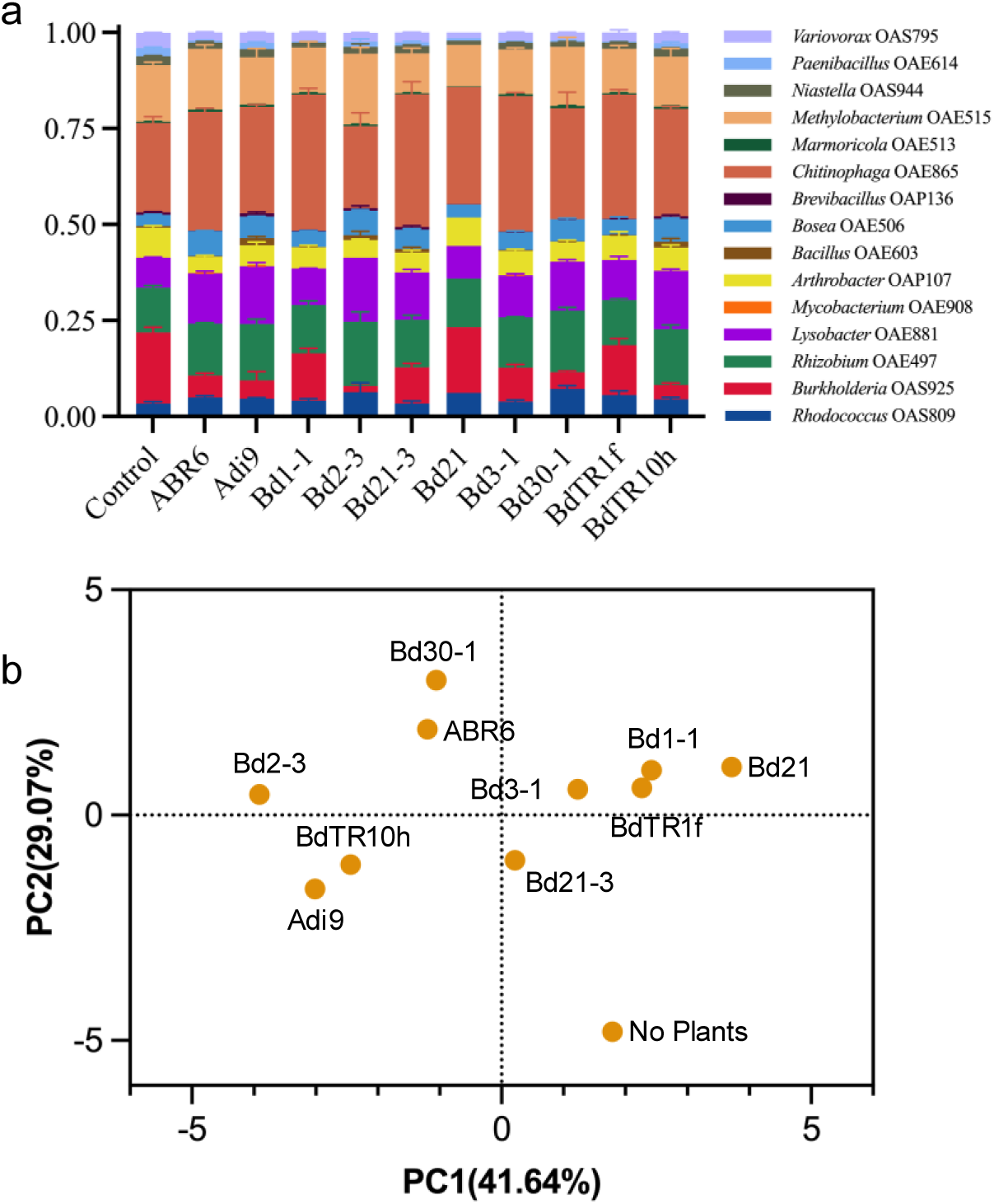
Distinct microbiome profiles across the ten *B. distachyon* lines. **a** Relative abundance of the SynCom members in the plant spent medium. Bacterial pellets were collected by centrifuging the plant medium for 16S rRNA sequencing; **b** Principal coordinate analysis showing the difference in the SynCom composition in the plant spent medium of ten *Brachypodium* lines. Averages of relative abundance of the SynCom members were used for the principal coordinate analysis. The SynCom inoculant was also added to the vessels filled with a plant-spent medium without plants (No Plants).

To explore chemical variation in root exudates of these *B. distachyon* lines, non-targeted metabolite profiling was conducted using liquid chromatography with tandem mass spectrometry (LC-MS/MS)^27^. In total, 3,644 metabolic features were detected (Supplementary Table 3). The nonmetric multidimensional scaling ordination (NMDS) plot suggests distinct metabolite abundance levels in root exudates of these *B. distachyon* lines in the absence of the SynCom (Supplementary Fig. 4). In comparison, the exudate metabolite profiles were less diverse in the presence of the SynCom (Supplementary Fig. 4), indicating that the SynCom presence buffered against differences in metabolite composition between *B. distachyon* lines that occurred in the absence of microbial partners.

To better understand the observed metabolic variation, we compared the similarities and differences in metabolic features detected in root exudates of these *B. distachyon* lines in the absence of the SynCom. Features detected in two or more lines account for 96 % of total features while only 4% were found in all lines (Supplementary Fig. 5a). We also found that the features unique to an individual line accounted for 18% of all features; for each individual line, the number varied between 6% to 12% of all features (Supplementary Fig. 5b). Collectively, these results demonstrate that these *B. distachyon* lines have distinct metabolic profiles in their root exudates in the absence of the SynCom, providing evidence for the distinct microbiome profile observed in the plant spent medium.

Metabolic feature changes were also examined in the root exudates of individual lines in response to the SynCom treatment. The NMDS ordination plot revealed that the inoculated plants formed a separate group (Supplementary Fig. 4). We also observed that the patterns of metabolic signal changes following the SynCom treatment were significantly different across these ten *B. distachyon* lines (Supplementary Fig. 6). Upregulated features varied from 6% to 27% of all features, while downregulated features ranged from 2% to 18% (Supplementary Fig. 6). Among these lines, BdTR1f, Bd21, and Bd1-1 had fewer features upregulated by the SynCom (6%, 6%, and 8%, respectively), whereas Bd30-1, BdTR1f, Bd2-3, and Bd21-3 had only 1%, 2%, 3%, and 5% of features downregulated, respectively (Supplementary Fig. 6). Overall, we did not observe any upregulated or downregulated features shared across all ten lines (Supplementary Fig. 7). Taken together, the metabolic changes in the root exudates in response to the SynCom varied across these *B. distachyon* genetic lines, providing further evidence for the observed distinct microbiome profile.

### Dopamine is a key dominant exometabolite modulating the SynCom composition and plant growth

Previously, we observed that dopamine was present in the spent medium of *B. distachyon*^27^. In this study, we also detected dopamine in the rhizosphere soil of *B. distachyon* (Supplementary Fig. 8). Interestingly, we found that the dopamine peak is one of the most intense peaks detected in both the *B. distachyon* root and the root exudates (Fig. 2a). Additionally, we observed natural variation in the dopamine levels in root exudates of the ten *B. distachyon* lines (Fig. 2b, 2c, Supplementary Table 4). In humans, dopamine is known to play a role in regulating gut microbe growth^21^, suggesting a possible role for dopamine in regulating rhizosphere microbial assembly.

**Fig. 2.**
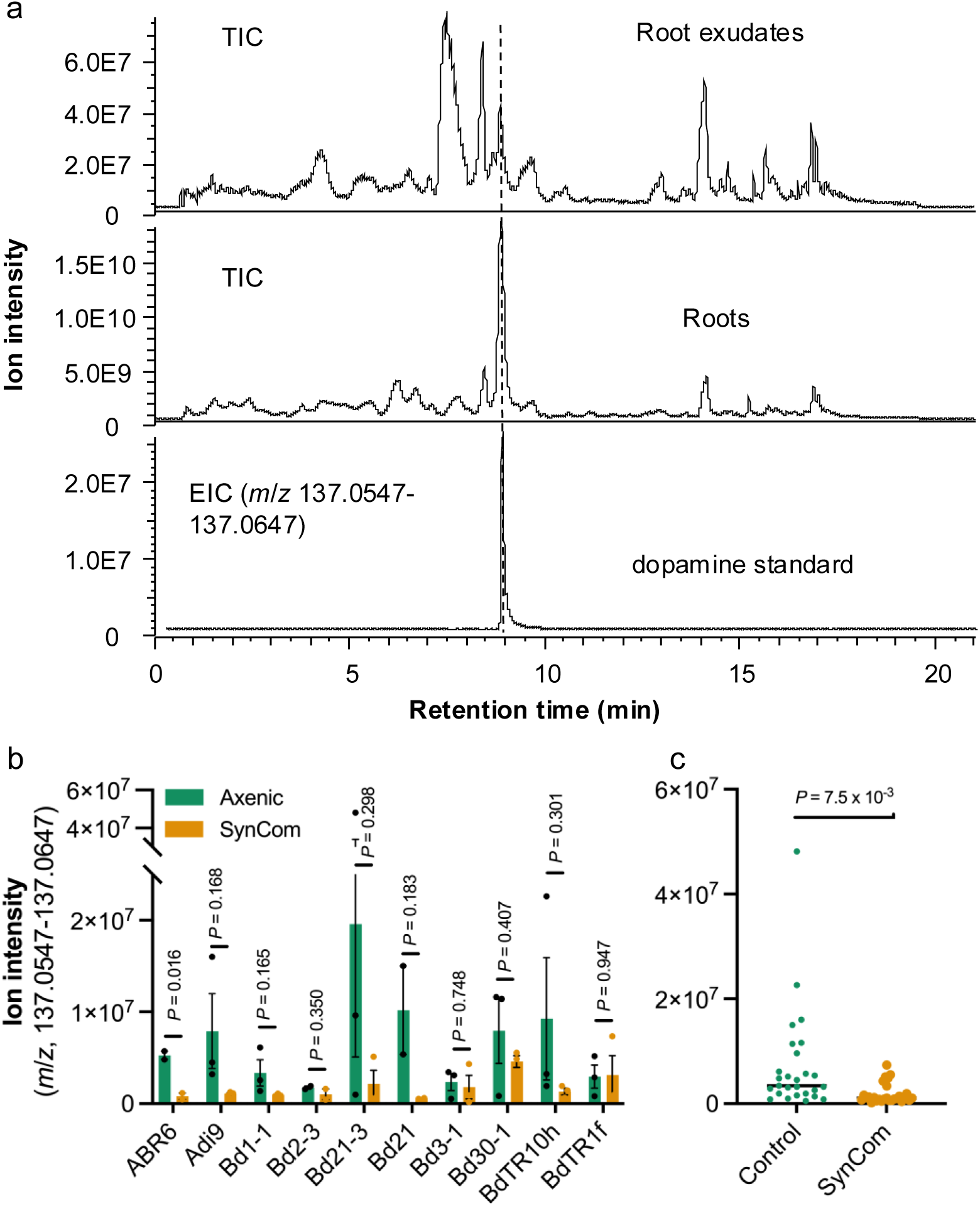
Dopamine is abundant in *B. distachyon* root exudates. **a** Representative LC-MS/MS TICs (total ion chromatograms) show dopamine as one of the most abundant metabolites in both *B. distachyon* exudates and roots. EIC stands for extracted ion chromatogram. *B. distachyon* Bd21-3 plants were grown in a hydroponic system. Plant spent media and the roots were collected from 4-week-old plants for LC-MS/MS analysis. **b** shows that dopamine levels vary across ten *B. distachyon* lines in the absence and presence of the SynCom. Ten *B. distachyon* lines were grown in a hydroponic system in the absence and presence of a 16-member bacterial SynCom. Plant spent media were collected for LC-MS/MS analysis at week 4. Error bars represent the mean ± SEM of the peak height (*m*/*z*, 137.0547-137.0647) (*n* = 2 or 3). **c** shows a combined analysis (*n* = 28 or 29) in the absence and presence of the SynCom. Overall, the SynCom negatively affected the dopamine levels in root exudates. The *P-*value was calculated using the student’s *t-*test.

To explore the possible function of dopamine in modulating plant-microbiome interactions, we performed an association analysis using the paired metabolomics and 16S rRNA amplicon sequencing data and found that dopamine levels in root exudates were significantly associated with the abundance of six SynCom members in the plant spent medium of the ten *B. distachyon* lines (Fig. 3a). Dopamine was positively associated with the relative abundances of two Actinobacteria, *Rhodococcus* OAS809 and *Marmoricola* OAE513, and three Proteobacteria, *Rhizobium* OAE497, *Lysobacter* OAE881, and *Bosea* OAE506, whereas it was negatively associated with another Proteobacteria, *Burkholderia* OAS925 (Fig. 3a).

**Fig. 3.**
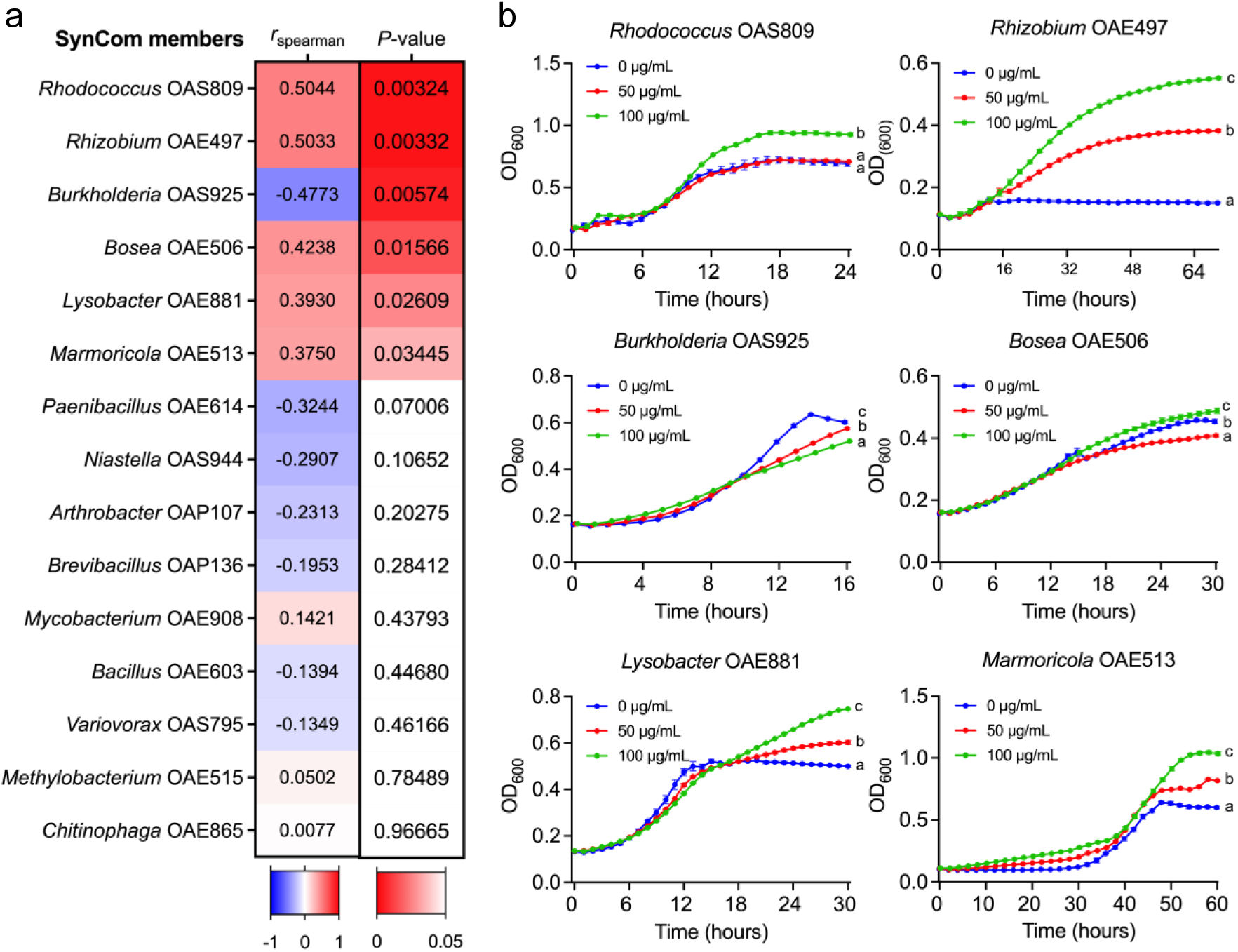
Dopamine regulates the SynCom composition in a hydroponic system. **a** Spearman correlation (*n* = 28) showing that dopamine levels were significantly associated with the relative abundance of six SynCom members either positively or negatively (*P* < 0.05, highlighted in red). **b** *In vitro* bacterial growth assays showing that dopamine regulates microbial growth. Average (*n* = 4; ± SEM) bacterial growth estimates (OD_600_) of six SynCom members in liquid media (R2A or 1/10 R2A) in the presence of dopamine at 0 (blue), 50 (red), and 100 (green) µg/mL. Within plots, different letters represent significant differences (one-way ANOVA, Tukey’s test corrections for multiple comparisons, *P* < 0.05).

To further investigate the mechanism underlying the potential role of dopamine in regulating the SynCom composition, we next performed *in vitro* assays to examine its growth effect on the six bacterial isolates whose abundances were significantly associated with the dopamine levels in the plant spent medium. To determine physiologically relevant concentrations for the assays, we measured dopamine concentration inside the roots since accurately estimating dopamine levels in root exudates is challenging. Based on an internal calibration curve of a [ring-^13^C_6_]-labeled dopamine (Supplementary Fig. 9a), we estimated that dopamine concentration could reach up to 2.71 ± 0.18 mg/g fresh weight in *B. distachyon* roots (Supplementary Fig. 9b). Next, we grew these six bacteria individually in liquid culture media supplemented with dopamine at two physiologically relevant concentrations and measured bacterial growth based on optical density at 600 nm (OD_600_) over time (Fig. 3b). Consistent with our association analysis, we observed that dopamine at 100 μg/mL significantly stimulated the growth of all five isolates whose abundances were positively associated with dopamine while inhibiting the growth of *Burkholderia* OAS925, whose abundance was negatively correlated with dopamine (Fig. 3b). These results demonstrate that dopamine may play an important role in modulating microbial abundance within a rhizosphere community context, likely by impacting microbe growth.

Interestingly, we also observed that dopamine levels in exudates were positively associated with plant morphological phenotypes in the hydroponic system, including root biomass, shoot biomass, root length, and total biomass (Supplementary Fig. 10a). Plant growth assays confirmed that dopamine was able to promote the growth of hydroponically grown *B. distachyon* plants at the two lower levels used in our experiment (Supplementary Fig. 10b). Our result is consistent with previous studies showing that dopamine can promote plant growth under various stress conditions^26^.

### Exogenous application of dopamine alters soil microbiota

Given that dopamine is correlated with the SynCom composition and regulates soil bacterial growth *in vitro*, we hypothesized that dopamine likely impacts soil microbial communities in a native context. Since the genes involved in dopamine biosynthesis remain unknown in plants, we elected to test this by adding dopamine in dry form into 20 g of agricultural soil at two physiologically relevant concentrations (1.5 mg and 3.0 mg per 20 g of dry soil) and maintaining the soil with a water content of approximately 24.5%. We also included norepinephrine as a control for carbon (C) and nitrogen (N) source input into the soil since norepinephrine is a direct derivative of dopamine and has the same C/N ratio; however, norepinephrine was only found at a trace level in *B. distachyon* roots as compared to the level of dopamine (Supplementary Fig. 11). The treatment was repeated twice a week over a period of 6 weeks. Based on sequencing analysis of the V4/V5 region of the 16S rRNA gene, zero-radius operational taxonomic units (zOTUs) were used to analyze soil bacterial diversity and membership across the soil treatments (Supplementary Table 5). Principal coordinate analysis revealed that both the chemical types and their concentrations were drivers in shaping the soil microbiome (Fig. 4a). The soil samples treated with the high dose of dopamine and norepinephrine formed two separate groups (Fig. 4a, Supplementary Fig. 12), suggesting that dopamine and norepinephrine at the high dose significantly, but differently altered the soil microbiota (Supplementary Fig. 12).

**Fig. 4.**
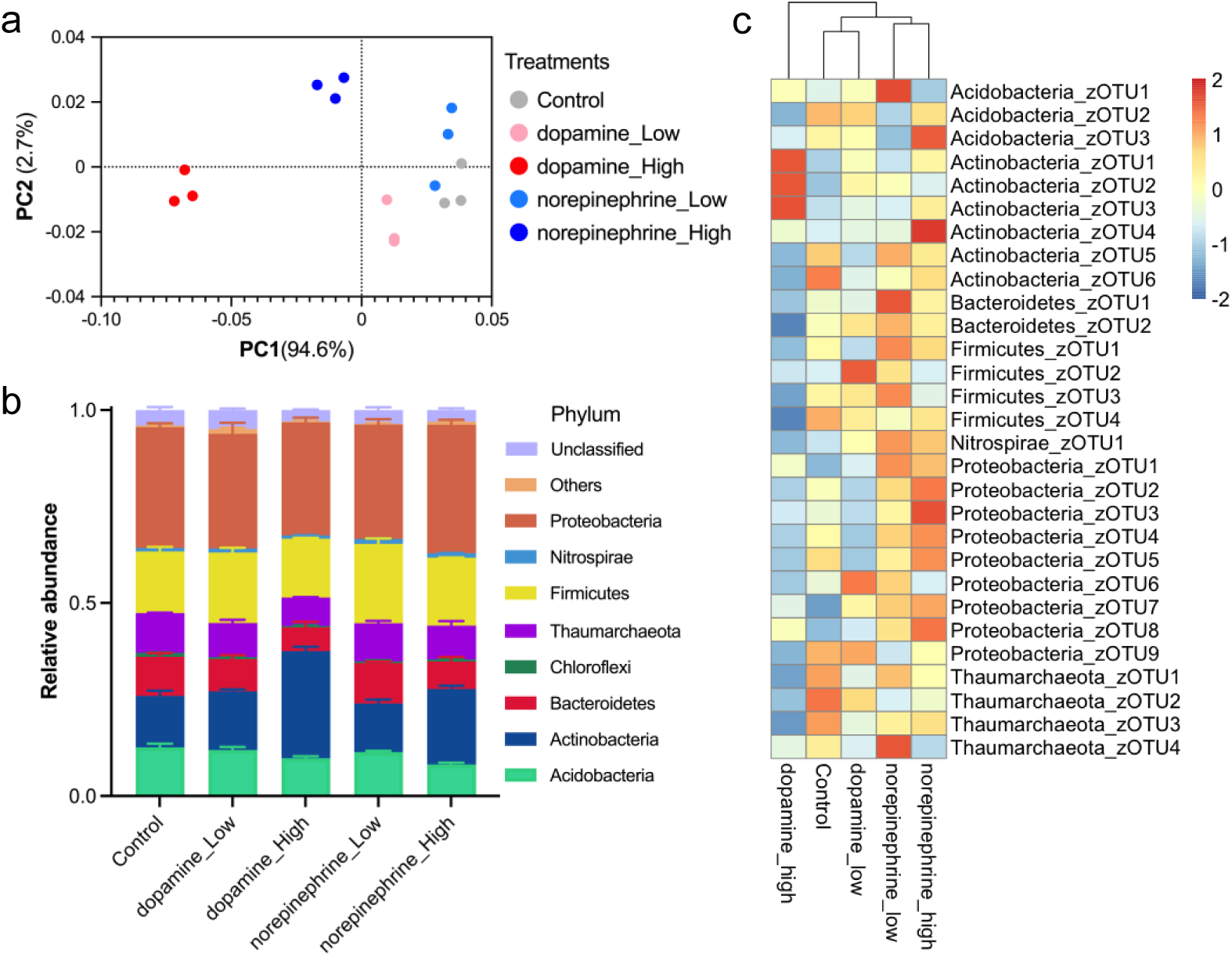
Dopamine modulates the soil microbiome. **a** The Principal coordinate analysis plot showing differences in bacterial community structures between samples. The color code depicts the different treatments (*n*=3). **b** Relative abundance of bacterial phyla in the soil of the control or treatment with plant metabolites, dopamine and norepinephrine. Low abundant phyla with <1% of the total reads in all samples are summarized as “Others”. Error bars represent mean ± SEM (*n* = 3). **c** Heatmap showing differences in relative abundances of bacterial zOTUs in soil after application of different plant metabolites at two concentrations. The heatmap was generated using a pheatmap package in R 4.4.2. The values scaled in the row direction are displayed in color ranging from blue (low) to red (high). Only zOTUs with a relative abundance ≥ 1% under at least one condition are shown. For a full list of zOTUs, see Supplementary Table 5.

Taxonomic analysis revealed that in the control soil samples, six phyla, Proteobacteria, Firmicutes, Actinobacteria, Acidobacteria, Thaumarchaeota, and Bacteroidetes, were the most abundant, with average relative abundances of 31.3%, 16.0%, 13.5%, 12.5%, 10.2%, and 10.2%, respectively, while others were less abundant (all < 2%) (Fig. 4b, Supplementary Fig. 13). Among all bacterial phyla, only Actinobacteria were significantly enriched in soils treated with both dopamine and norepinephrine at the high dose; however, dopamine had a greater effect than norepinephrine (Fig. 4b, Supplementary Fig. 13). Additionally, we observed that Acidobacteria significantly decreased in the soil treated with the high dose of norepinephrine as compared to those in the control soil, but there was no significant difference in the relative abundance of Acidobacteria between treatments with dopamine and norepinephrine (Supplementary Fig.13). These results underscore the selective impact of these metabolites, particularly dopamine, on soil Actinobacteria communities.

The treatment with dopamine and norepinephrine also resulted in characteristic changes at the zOTU level within multiple bacterial phyla (Fig. 4c, Supplementary Fig. 14a, b). With principal component analysis, a notable trend was detected among zOTUs with relative abundance ≥0.1%, represented by principal component 1 (PC1). The responses of zOTUs to treatments were divided into two groups: the first group included dopamine (both low and high) and norepinephrine_High, while the second group consisted of the Control and norepinephrine_Low (Supplementary Fig. 14a, b). Further statistical analysis identified eight sensitive zOTUs that statistically (|loading score| ≥ 0.10) contributed to this trend, and some of them were differentially enriched among treatments (*P*-value < 0.05) (Fig. 4c, Supplementary Fig. 14b), suggesting that these metabolites had a specific role in modulating the soil microbial communities. The relative abundances of Actinobacteria_zOTU1 and Actinobacteria_zOTU3 were significantly increased by both dopamine_High and norepinephrine_High (Fig. 4c, Supplementary Fig. 14b). Additionally, Actinobacteria_zOTU2 and Actinobacteria_zOTU4 were enriched by dopamine_High and norepinephrine_High, respectively. Conversely, Proteobacteria*_*zOTU4, Bacteriodetes_zOTU1, Thaumarchaeota_zOTU1, and Actinobacteria_zOTU7 were more abundant in the Control and low norepinephrine treatments and were negatively impacted by high dopamine (Supplementary Fig. 14a, b). Taken together, these results suggest that treatment with these root-derived metabolites, particularly dopamine, specifically altered the bacterial community structures in the soil.

## Discussion

Root exudates play a crucial role in mediating interactions between plants and soil microbiota^8,30^. Genetic variation in root exudation is a key factor in the adaptability and ecological success of plants, influencing both individual plant growth and broader ecosystem dynamics^31,32^. In this study, we investigated the natural variation in root exudates of diverse *B. distachyon* lines and their influence on the composition of a synthetic microbial community in a hydroponic system. Our findings demonstrate the significant impact of intraspecific root exudate variation on shaping the composition and activity of a synthetic microbial community. The observed differences in exudate metabolic profiles across diverse *Brachypodium* lines underscore the role of genetic variation in governing root exudate composition, which subsequently influences microbial recruitment and community assembly, thereby contributing to the establishment of distinct root microbiome profiles. Through association analysis of paired exudate metabolomics and 16S rRNA sequencing data, we identified a dominant root exometabolite, dopamine, which significantly impacts soil microbial communities. Our results align with previous studies demonstrating the importance of plant genetic variation in modulating root exudate composition and subsequently affecting microbiome dynamics^33,34,35^. Understanding the natural variation in root exometabolites and its link to microbiome structure provides valuable insights into plant-microbiome interactions and can guide strategies for harnessing microbial consortia to enhance plant health and productivity in agricultural systems.

Integrative analysis of paired microbiome-metabolome data sets has proven to be the most promising strategy for the discovery of microbe-metabolite links in the human gut^36,37^. In this study, we demonstrate the utility of integrative analysis for uncovering microbe-metabolite links in the model grass *B. distachyon*. We first incubated diverse *Brachypodium* lines with a 16-member SynCom in a hydroponic system and collected paired exudate metabolomic data and 16S rRNA sequencing data. Using this data, we identified a dominant exometabolite, dopamine, in exudates that significantly associated with the abundance of six SynCom members (Fig. 3). The use of a hydroponic system in this study was crucial because it provided a controlled environment, facilitated exudate collection, and ensured adequate growth and sampling opportunities for the SynCom members^12,13,14^, enabling robust associations between metabolites and microbes. Though there are limitations in the correlation-based analysis to identify key microbiome-metabolite links, statistically significant correlations can be valuable for generating hypotheses and directing experimental research efforts. To confirm the role of dopamine in plant-microbiome interactions, we performed *in vitro* bacterial growth and soil treatment assays (Fig.3 & 4), both of which support the importance of dopamine in modulating soil microbial communities.

Consistent with previous studies in humans^20^, here we observed differential responses of selected soil bacteria to dopamine *in vitro*. Functional tests in soil revealed that catecholamines, particularly dopamine, could restructure the soil microbiota by specifically enriching Actinobacteria populations, underscoring the selective influence of dopamine on rhizosphere microbial communities. The underlying mechanism of dopamine modulating rhizosphere microbiota could be similar to that involved in regulating human gut microbial growth, such as affecting gene expression and biofilm formation^23,35^; however, it still needs further investigation.

Actinobacteria are known for their diverse metabolic capabilities and beneficial roles in plant health. Previous studies revealed the enrichment of Actinobacteria in drought-treated soils across various environments^38,39^, within drought-affected rhizospheres^40,41^, and within the host roots^42^, suggesting that Actinobacteria play an important role in plant stress response. Recently, two Actinobacteria (*Pseudarthrobacter* sp. L1D14 and *Pseudarthrobacter picheli* L1D33) were reported to promote plant growth in a hydroponic system^43^. This study extends these findings by demonstrating dopamine-mediated enrichment of Actinobacteria in both hydroponic and soil environments, suggesting a potential mechanism for plants to modulate soil microbial communities. Future investigations should focus on elucidating the molecular mechanisms underlying dopamine’s effects on microbial gene expression and community dynamics as well as regulation of dopamine production and exudation in plant hosts. Such insights could inform strategies for harnessing root exudate-derived metabolites to optimize plant-microbe interactions in agricultural systems, promoting sustainable soil management practices and enhancing crop productivity.

## Materials and methods

### Plant materials and plant growth

Ten inbred *B. distachyon* lines (Adi9, ABR6, Bd1-1, Bd2-3, Bd21, Bd21-3, Bd3-1, Bd30-1, BdTR10h, and BdTR1f) were used in this study^44,45,46,47^. Seeds were surface sterilized in 70% (v/v) ethanol for 30 s and 5 min in 6% (w/v) sodium hypochlorite, followed by five washes with sterile water. The seeds were subsequently stratified on 1.5% (w/v) agar plates containing 0.5 × Murashige & Skoog (0.5 × MS, MSP01, Caisson Laboratories, USA) and placed in darkness at 4°C for three days. The plates containing the seeds were then moved to a growth chamber with a temperature of 22°C, photosynthetic photon flux density at 150 μmol m^−2^s^−1^, and a 16 h light/8 h dark photoperiod for two days.

The culture vessels (PTL-100™, PhytoTech Labs) were rinsed five times with MilliQ water and autoclaved. Two seedlings of *B. distachyon* were placed onto a floating holder and then transferred to each vessel with 40 mL of 0.5 × MS. Plants were grown in a 16 h light/8 h dark regime at 22 °C and 50% relative humidity with 150 μmolm^-2^s^-1^ illumination for four weeks. The culture vessels without plants filled with the growth medium were incubated in the same conditions as the controls. For plant growth with dopamine (Sigma-Aldrich), seven-day-old *B. distachyon* seedlings were treated with dopamine at concentrations of 25, 50, and 100 μM for three weeks. At week 4, *B. distachyon* plants were harvested. Root and shoot fresh biomass were measured. Root length and shoot length were quantified using the SmartRoot plugin (version 4.21) in ImageJ (version 2.0.0). The sterility of the hydroponic setup was examined before exudate collection by plating 50 μL of medium on Luria-Bertani (LB) plates, followed by seven-day incubation at 30 °C.

### Bacterial strains and synthetic community inoculation

Sixteen bacterial strains used in this study were isolated from the rhizosphere and bulk soil surrounding a *Panicum virgatum* (switchgrass) plant in Oklahoma, United States^29^. Strains were cultured at 30°C and 200 rpm in the following liquid media: *Lysobacter* OAE881, *Burkholderia* OAS925, *Variovorax* OAS795, *Chitinophaga* OAE865, *Bosea* OAE506, *Rhodococcus* OAS809, *Marmoricola* OAE513, *Paenibacillus* OAE614, *Methylobacterium* OAE515, *Arthrobacter* OAP107, *Mucilaginibacter* OAE612, and *Brevibacillus* OAP136 in Reasoner’s 2A (R2A) medium^48,49^, *Niastella* OAS944, *Rhizobium* OAE497, and *Mycobacterium* OAE908 in 0.1 × R2A, and *Bacillus* OAE603 in Luria-Bertani (LB) medium (Thermo Fisher, USA) (Supplementary Table 1).

Bacterial cultures were centrifuged at 3,600 g for 20 minutes and washed 3 times with 0.5× MS. Individual bacterial strains were resuspended to have an OD_600_ at 1.0, and then equal volumes were pooled together to make the 16-member SynCom. The SynCom inoculant (800 μL) was applied to the 0.5× MS growth medium of half of the culture vessels containing *B. distachyon* (n = *2* or *3*) at week one to have a final OD_600_ of approximately 0.02. The SynCom inoculant was also added to the vessels filled with plant-spent medium without plants (No Plants). Following inoculation, plants were grown for an additional 3 weeks under a 16 h light/8 h dark cycle, 22°C, and 50% humidity until harvest.

### Exudate harvesting and bacterial cell collection

Four-week-old plants were carefully removed from the culture vessel together with the floating holder, and the plant spent medium was collected into a 50 mL falcon tube. OD_600_ was measured to determine the SynCom growth. The plant-spent medium was then centrifuged at 3,600 g for 20 minutes to collect bacterial cells. The supernatants were filtered with 0.2 μm PES membrane filters (Pall Corporation, NY, USA), freeze-dried (Labconco Freeze-Zone), and stored at -80 °C for metabolite analysis. The bacterial pellets were resuspended in 20% glycerol and stored at -80 °C prior to DNA extraction.

### LC-MS/MS analyses of *B. distachyon* root exudates

The plant spent media from 4-week-old plants (approximately 30 mL) were frozen at -80°C and subsequently freeze-dried using a Labconco Freeze-Zone. The dried material was resuspended by rinsing the tubes with 1 mL of LC-MS grade methanol (Sigma-Aldrich), which was then transferred to 2-mL tubes and dried in a Thermo Speed-Vac concentrator. The dried material was then resuspended in 300 μL of LC-MS grade methanol containing internal standards. The solution was vortexed for 2 × 10 seconds, bath-sonicated in ice water for 15 minutes, and centrifuged at 10,000 g for 5 minutes at 10°C to pellet the insoluble materials. The supernatants were then filtered using 0.22-μm polyvinylidene difluoride microcentrifuge filtration devices (Pall Corporation, NY, USA) at 10,000g for 5 minutes at 10°C. The filtrates were used for metabolite analysis. Polar metabolites were separated using hydrophilic interaction chromatography (HILIC) and detected on a Thermo Q Exactive Hybrid Quadrupole-Orbitrap Mass Spectrometer. LC-MS/MS. Briefly, an InfinityLab Poroshell 120 HILIC-Z, 2.1×150 mm, 2.7 um column equipped on an Agilent 1290 HPLC stack was used for separations. Data was collected using data-dependent MS2 acquisition to select the top two most intense ions not previously fragmented within 7 seconds. Internal and external standards were used for quality control purposes. Method parameters are defined in Supplementary Table 6.

### Untargeted metabolomics analysis

The MZmine2 version 2.39 workflow was used for picking up features for positive and negative polarities^56^. A baseline filter was initially performed, which accepted features with a retention time (RT) of >0.6 min, a maximum peak height of >1 x 10^6^, and >10 x the maximal peak height in extraction (ExCtrl, *n* = 3) and technical controls (TeCtrl, *n* = 3). To remove the metabolites from the SynCom bacterial members, features were further filtered to remove those with a maximum peak height of >1 x 10^6^ detected in No Plants controls which were filled with a root spent medium and the SynCom. We added +0.1 to all filtered features during the fold change calculations to avoid errors when dividing by zero. These parameters were used for feature filtering for all *Brachypodium* lines in the presence and absence of the SynCom.

### Association analysis of dopamine with the SynCom members and plant phenotypes

To determine the covariance between dopamine and the composition of the microbial communities in the plant-spent media, we performed association analysis based on the relative levels of dopamine (ion intensity of 137.0547-137.0647 *m*/*z*) and the relative abundances of SynCom members in the plant-spent medium of the ten *B. distachyon* lines. Spearman correlations and *P*-values were calculated using the *rcorr* function of the R package *Hmisc* for dopamine-SynCom member pairs. Positive and negative correlations correspond to positive and negative links, respectively. Additionally, Spearman correlations and *P*-values were also calculated for dopamine-phenotype pairs.

### Soil treatment

Soil (Yolo silt loam) was collected from an agricultural field at UC Davis (38°32’16.9“N, 121°46’00.2”W). After collection, the soil was dried at room temperature and stored at 4°C for 2-3 months, and then sieved (2 mm mesh) before starting the experiment. The metabolites (dopamine and norepinephrine) at two doses (1.5 mg and 3.0 mg) were directly mixed into 20 g of dried soil, and then 6.5 mL of water was added to achieve a water content of approximately 24.5% by weighing each replicate and adding water accordingly, and the treatments were noted with dopamine_Low, dopamine_High, norepinephrine_Low, and norepinephrine_High, respectively. The treatments were repeated twice a week over a period of 6 weeks, and the water content was maintained at approximately 24.5%. Control soil was watered without the addition of any metabolites. The soil was covered with foil and incubated at room temperature. Three replicates were prepared per treatment. Soil samples were harvested at week 6 and stored at -20°C prior to DNA extraction.

### DNA extraction, 16S rRNA amplicon generation, and sequencing

Total DNA was extracted from samples using the FastDNA SPIN Kit for Soil (MP Biomedicals, United States) as described in the manufacturer’s protocol. Genomic DNA was then eluted in 40 μL of nuclease-free water. The V4/V5 region of the 16S rRNA gene was amplified by PCR from each of the purified DNA samples using the 515F/926R primers based on those from the Earth Microbiome Project^50,51^, but with in-line dual Illumina indexes. The resulting amplicons were sequenced on an Illumina MiSeq using 600 bp v3 reagents. Custom Perl scripts were employed for read processing, which included merging with Pear^52^, filtering reads (zero-radius operational taxonomic units, zOTUs) with more than one expected error using Usearch^53^, demultiplexing using inline indexes, and filtering rare reads and chimeras with Unoise^54^. 16S sequences were determined by comparing them against the RDP database for taxonomy assignment^55^. zOTU is interchangeable with ESV (exact sequence variant) and ASV (amplicon sequence variant).

### Processing and statistical analysis of 16S rRNA counts

Phylum-level tables were generated using the taxonomic classifications performed above, with all zOTU counts assigned to a given phylum summed in the table. The denoised table of zOTU counts and phylum tables were trimmed to remove low abundance (i.e., zOTUs only present in 1 or 2 samples with fewer than 10 reads) and then analyzed using the DESeq2 software program with default settings. Treatment comparisons were performed using the contrast function.

### Quantification of dopamine in *Brachypodium* roots

*Brachypodium* Bd21-3 root tissues were harvested from 4-week-old hydroponically grown plants and stored at -80 °C for further analysis. Before extraction, root tissues (approximately 0.1g fresh weight) were ground into powder using bead milling. 500 μL extraction buffers [95% (70% methanol, methanol:H_2_O_2_, 70:30, v/v) and 5% 1 M hydrochloric acid] with a [ring-^13^C_6_]-labeled dopamine (Cambridge Isotope Laboratories) at concentrations of 5 mM, 10 mM, 20 mM, 40 mM, and 80 mM were then added into the root powders (*n* = 2) for dopamine extraction overnight at 4°C. The samples were subsequently centrifuged for 10 min at 15,000 g. The supernatants were further filtered using 0.22-μm polyvinylidene difluoride microcentrifuge filtration devices (Pall Corporation, NY, USA) by centrifuging (10,000 g for 5 min at 10°C) and diluted 100-fold before LC-MS/MS analysis using hydrophilic interaction chromatography (HILIC) on a Thermo Q Exactive Hybrid Quadrupole-Orbitrap Mass Spectrometer.

The dopamine calibration curve was obtained by analyzing the above samples on LC-MS/MS. The analytical curve was described by a linear regression model *y* = **a***x* + **b**, where *y* represents the analytical response, *x* is the concentration of analyte in μM, **a** denotes the slope of the curve, and **b** is the intercept. The calibration curve equation and the *R*^2^ determination were established based on the linear regression of the peak height of the [ring-^13^C_6_]-labeled dopamine (*m*/*z*, 160.1014-160.1114) over its concentration. The calibration curve (*y* = 5.69E5*x* + 4.26E5) showed good linearity in the range of 5.0-80.0 μM tested with the coefficient of determination *R*^2^= 0.9996 (Supplementary Fig. 9a). The levels of endogenous dopamine (*m*/*z*, 154.0812-154.0913) in *Brachypodium* roots were calculated based on the [ring-^13^C_6_]-labeled dopamine calibration curve (*n* = 12).

### *In vitro* Bacterial growth

*In vitro* bacterial growth assays with dopamine were performed using the Clinical and Laboratory Standards Institute M38-A2 guidelines^57^. In brief, a 96-well microtiter plate-based method using a Synergy4 (BioTech Instruments) reader was used to monitor bacterial growth at 30°C in liquid media through periodic measurements of changes in optical density (OD_600_ nm) for 72 h. Bacterial cultures were centrifuged at 3,600 g for 20 minutes and washed 3 times with the corresponding growth media. Each plate well contained 200 µL of initial bacterial inoculum (OD_600_ at approximately 0.01) supplemented with dopamine at two different concentrations (50 and 100 µg/mL). Methanol was used as a solvent for making dopamine solutions. Our controls were blank medium and bacterial-inoculated medium with 0.1% methanol (*n* = 4).

### Statistical analysis

Statistical analyses were conducted using JMP Pro v.13.0 (SAS Institute) and Prism v.9.0 (GraphPad). One-way ANOVA was performed to evaluate statistical differences. Tukey tests were used to correct for multiple comparisons between control and treatment groups. Student’s unpaired two-tailed *t*-tests were conducted for pairwise comparisons. A *P*-value of < 0.05 was considered to be statistically significant.

The beta-diversity of soil microbial communities was calculated as Bray-Curtis Dissimilarity using the *vegan* package, and permutational multivariate analysis of variance (PERMANOVA) was performed using the *adonis2* package to test their significant differences between treatments, both done in the R environment (version 4.4.1.). The sensitive zOTUs were selected using the modified selection pipeline^58^. Briefly, principal component analysis was performed on zOTUs with maximum relative abundance of at least 0.1%, after scaling their relative abundances using the package *zCompositions*. The principal component (PC) scores were analyzed with one-way ANOVA and PC with significant differences between treatments (*P*-value < 0.05) were selected. Then, zOTUs with an absolute value of the loading score larger than |0.1| were selected as sensitive taxa. The lsmeans of PC scores for each treatment were multiplied by the loading scores of each sensitive taxa to visualize their responses to the treatments. Then, hypothesis testing was performed on the relative abundances of these sensitive taxa by treatment using one-way ANOVA and post-hoc Tukey’s test. The figures for PERMANOVA and sensitive taxa were created using *ggplot2* in the R environment^59^.

## Acknowledgments

The author(s) declare financial support was received for the research, authorship, and/or publication of this article. YD and TN are supported by the m-CAFEs Microbial Community Analysis & Functional Evaluation in Soils, a Science Focus Area at Lawrence Berkeley National Laboratory funded by the U.S. Department of Energy, Office of Science, Office of Biological & Environmental Research DE-AC02-05CH11231, and an Award DE-SC0021234 led by UC San Diego from the U.S. Department of Energy, Office of Science, Office of Biological & Environmental Research.

## Author contributions

Y.D. designed the study with input from J.V.P. and T.R.N. Y.D. and Y.Z. performed the assays to investigate the genetic variation of root exudates in plant-microbiome interactions. Y.D. and H.V. conducted *in vitro* bacterial growth and soil treatment assays. Y.D., P.F.A., H.K.C., N.K., and D.C. analyzed the 16S rRNA sequencing data. Y.D., Y.Z., A.N.G., V.N., B.P.B., and S.M.K. collected and analyzed the metabolomics data. Y.D. wrote the manuscript with input from all the other authors. All authors read and approved the final manuscript.

---

**Supplementary Fig. 1.**
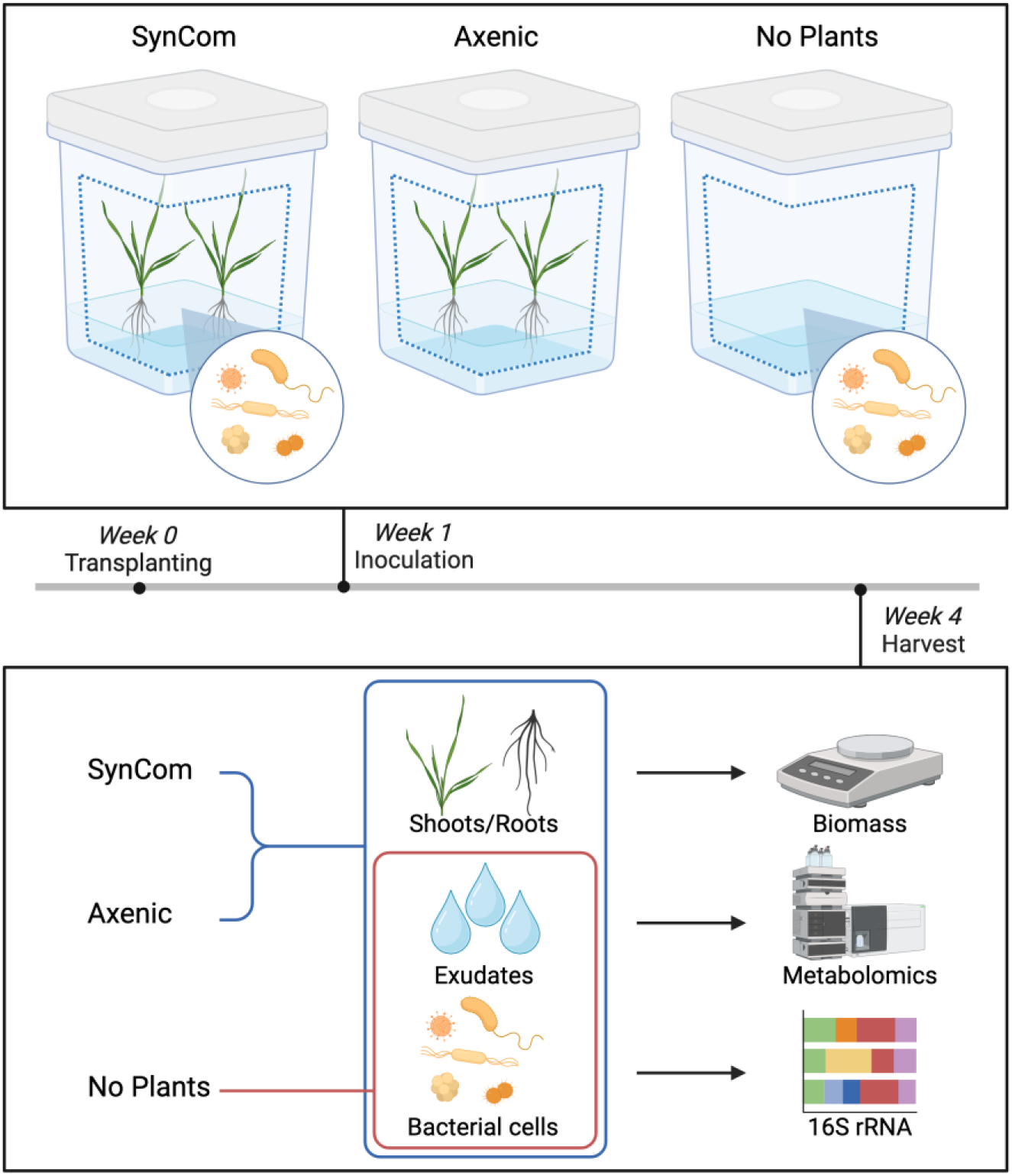
Schematic view of the experimental design for exploring the genetic variation of root exudates in plant-microbiome interactions. One-week-old hydroponically grown seedlings of ten *B. distachyon* lines were inoculated with a 16-member bacterial SynCom. Phenotypes were measured after an additional three-week incubation. Plants without the SynCom were used as the axenic controls. Exudates with the SynCom without plants were included (No Plants). We sampled the plant spent medium for exudate analysis by LC-MS/MS at week 4. The figure was created with <u>BioRender.com</u>.

**Supplementary Fig. 2.**
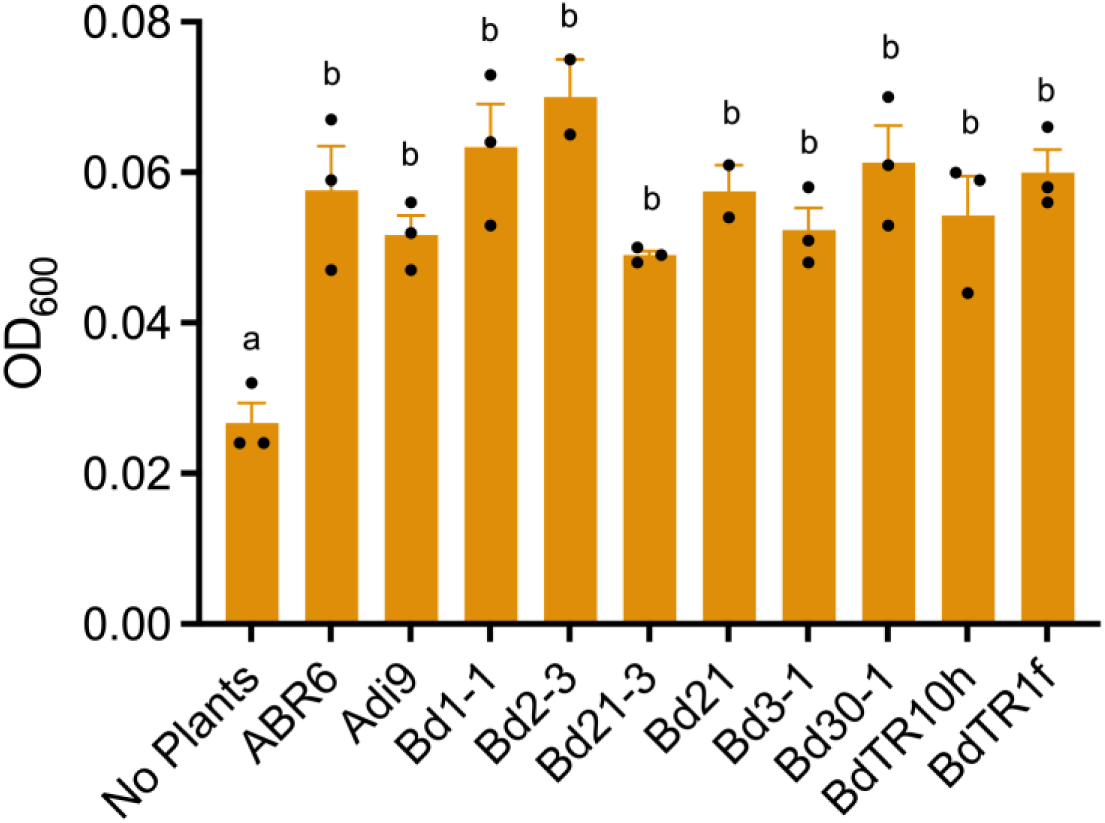
The growth of the SynCom bacteria in the plant spent medium as determined by the final OD_600_ measurement. Error bars represent mean ± SEM (*n* = 2 or 3). Within the plot, different letters (a–b) represent significant differences (one-way ANOVA followed by Tukey’s test corrections for multiple comparisons; *P* < 0.05). No Plants was used as the Control.

**Supplementary Fig. 3.**
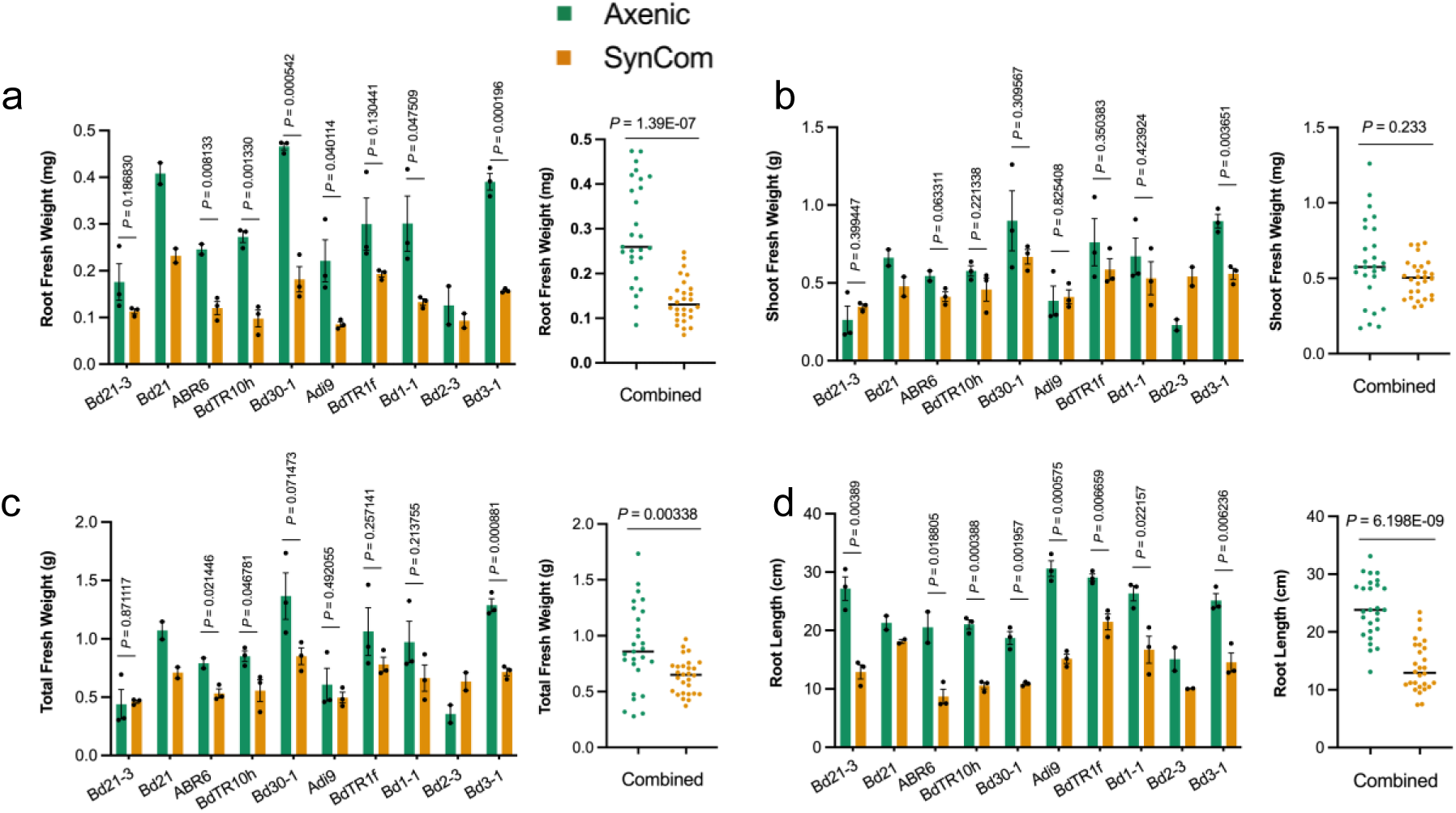
Phenotypic variation of ten *B. distachyon* lines in the absence and presence of a SynCom. Ten *B. distachyon* lines were grown in a hydroponic system for one week, and then half of them were inoculated with a 16-member bacterial SynCom. Phenotypes, including fresh root biomass (**a**), fresh shoot biomass (**b**), total fresh biomass (**c**), and root length (**d**), were measured after an additional three-week incubation. Error bars represent mean ± SEM (*n* = 2 or 3). Combined analyses of each phenotype in the absence and presence of the SynCom were also performed. The *P-*value was calculated using the student’s *t-*test.

**Supplementary Fig. 4.**
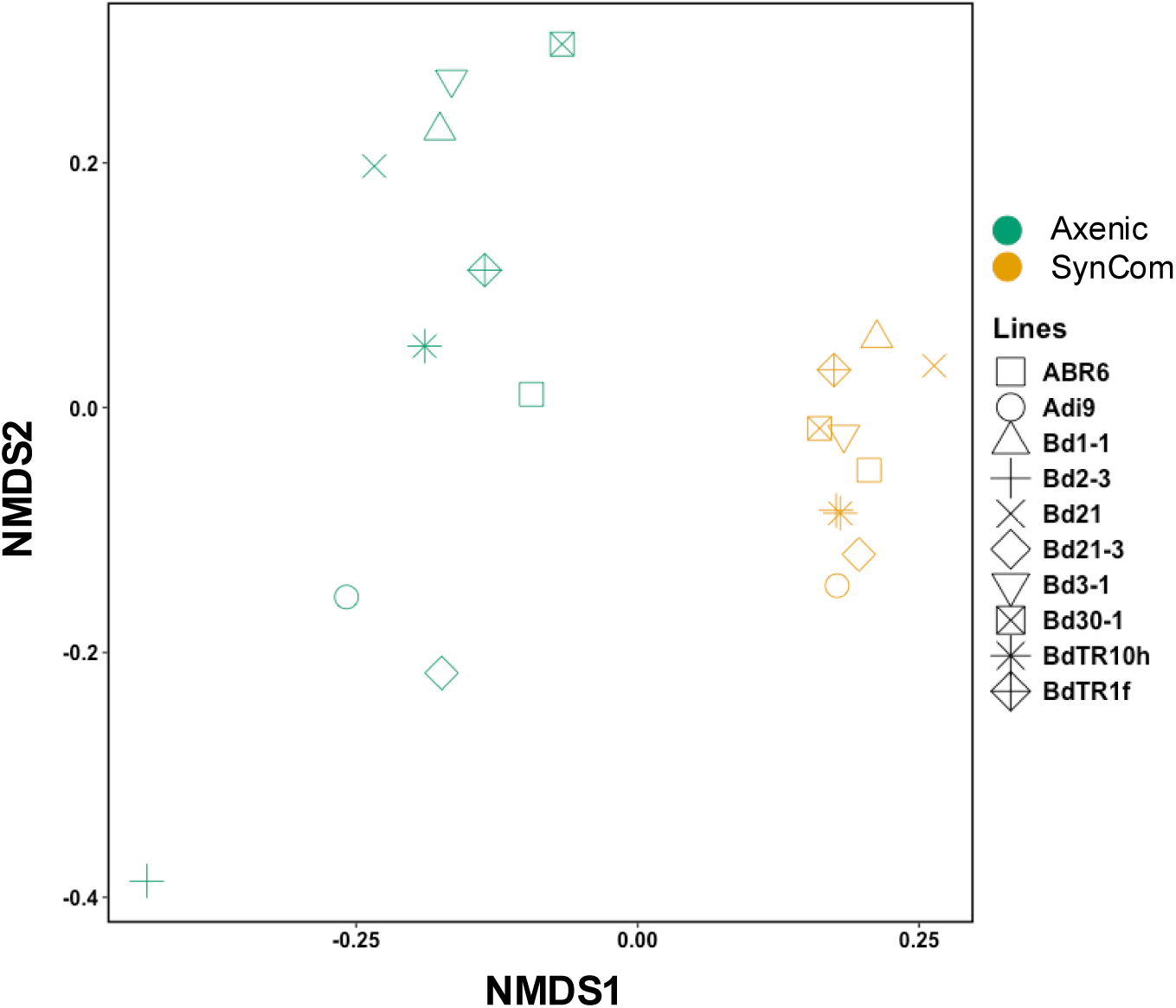
Non-metric multidimensional scaling (NMDS) plot for untargeted metabolomics analysis. The plot was constructed based on averaged peak heights (*n* =2 or 3) of features detected in the plant spent medium of ten *B. distachyon* lines in the absence and presence of the SynCom.

**Supplementary Fig. 5.**
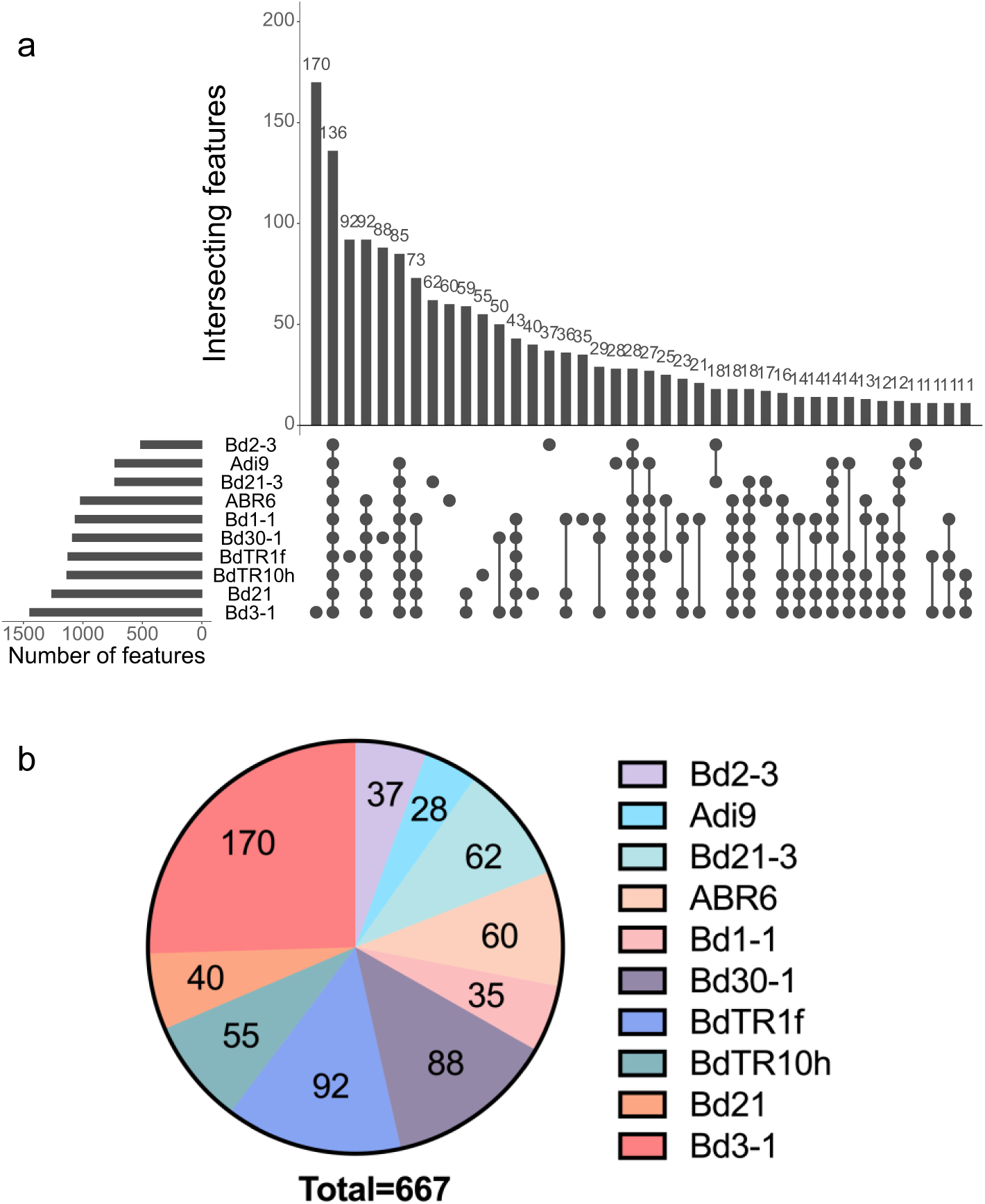
The number of features detected in root exudates. **a** The UpSet plot shows the number of shared filtered features detected in root exudates of ten *B. distachyon* lines in the absence of the SynCom. One hundred thirty six features (approximately 2%) are shared by the ten *Brachypodium* lines. **b** The pie chart shows the number of unique features detected in root exudates of individual *B. distachyon* lines in the absence of the SynCom.

**Supplementary Fig. 6.**
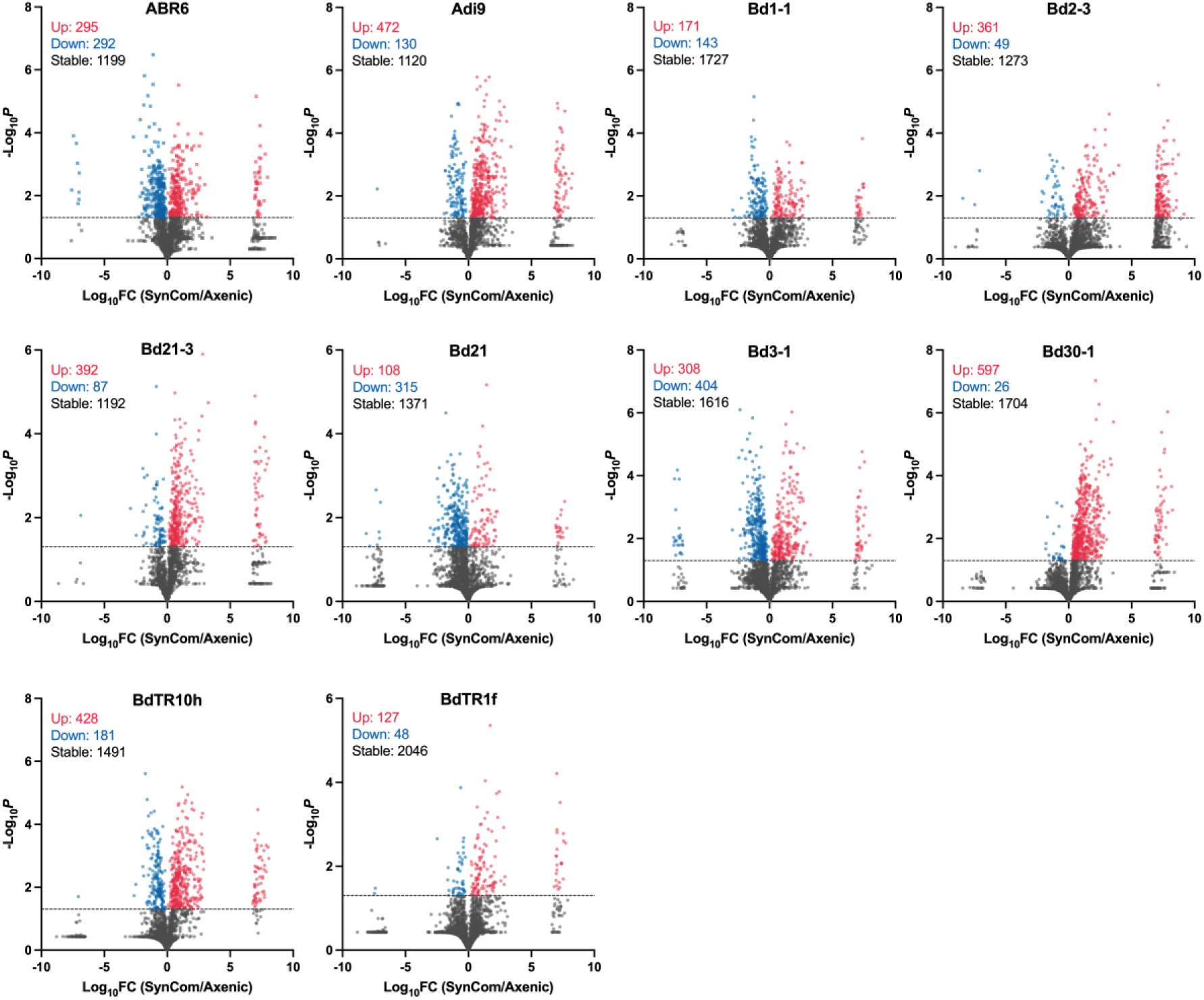
Exudate feature changes of individual *B. distachyon* lines in response to the SynCom. Volcano plots showing the Log_10_ fold changes (Log_10_FC) of features detected in root exudates of individual lines treated with the SynCom versus the axenic control (x-axis) and -Log_10_*P* values derived from student *t*-tests (y-axis). Statistically up-regulated features are labeled in red, and down-regulated features are labeled in blue (*P* < 0.05).

**Supplementary Fig. 7.**
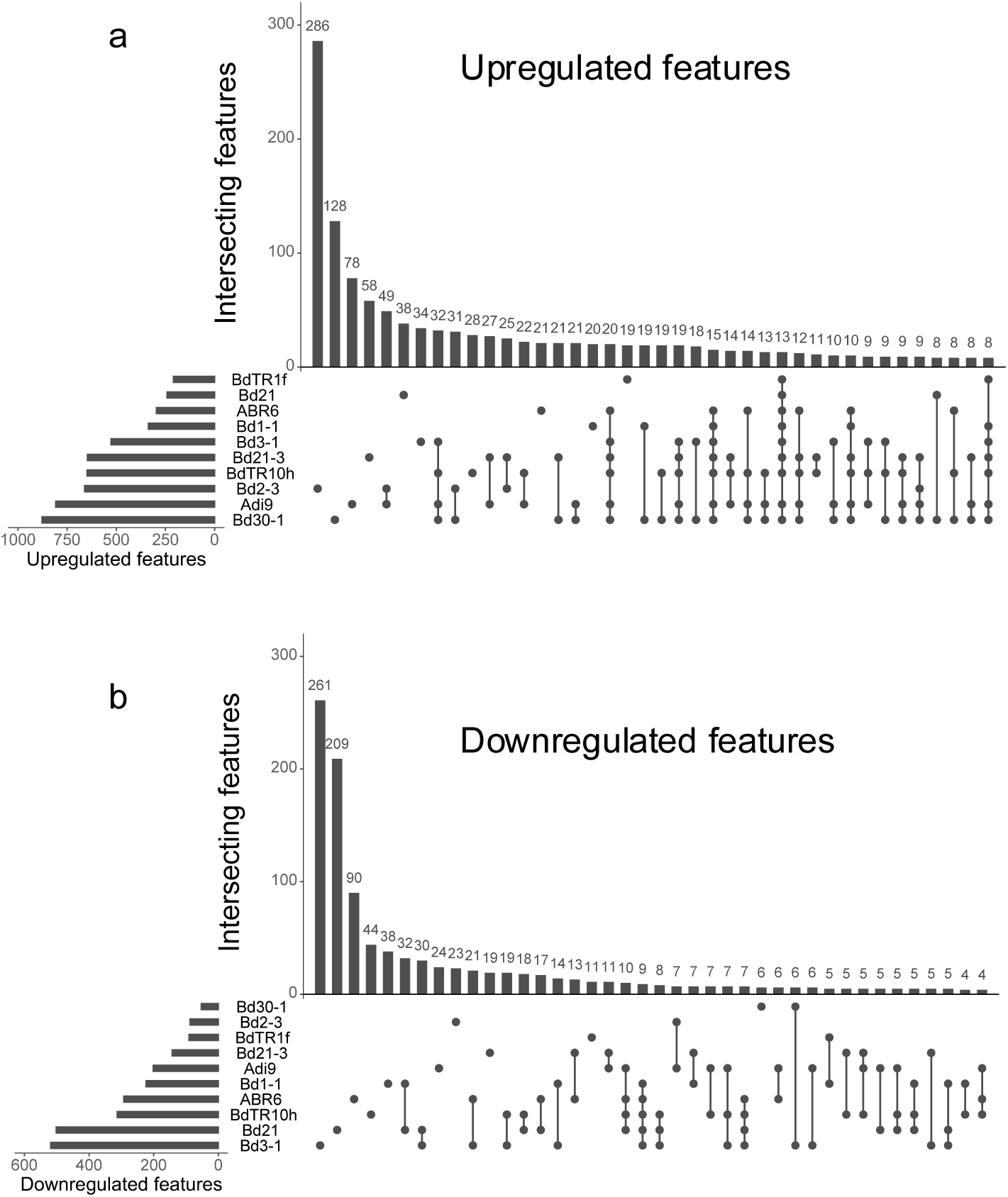
The number of significantly changed features in root exudates of ten *B. distachyon* lines. The UpSet plots show the numbers of shared features that are significantly upregulated (**a**) and downregulated (**b**) in root exudates of ten *B. distachyon* lines in response to the SynCom.

**Supplementary Fig. 8.**
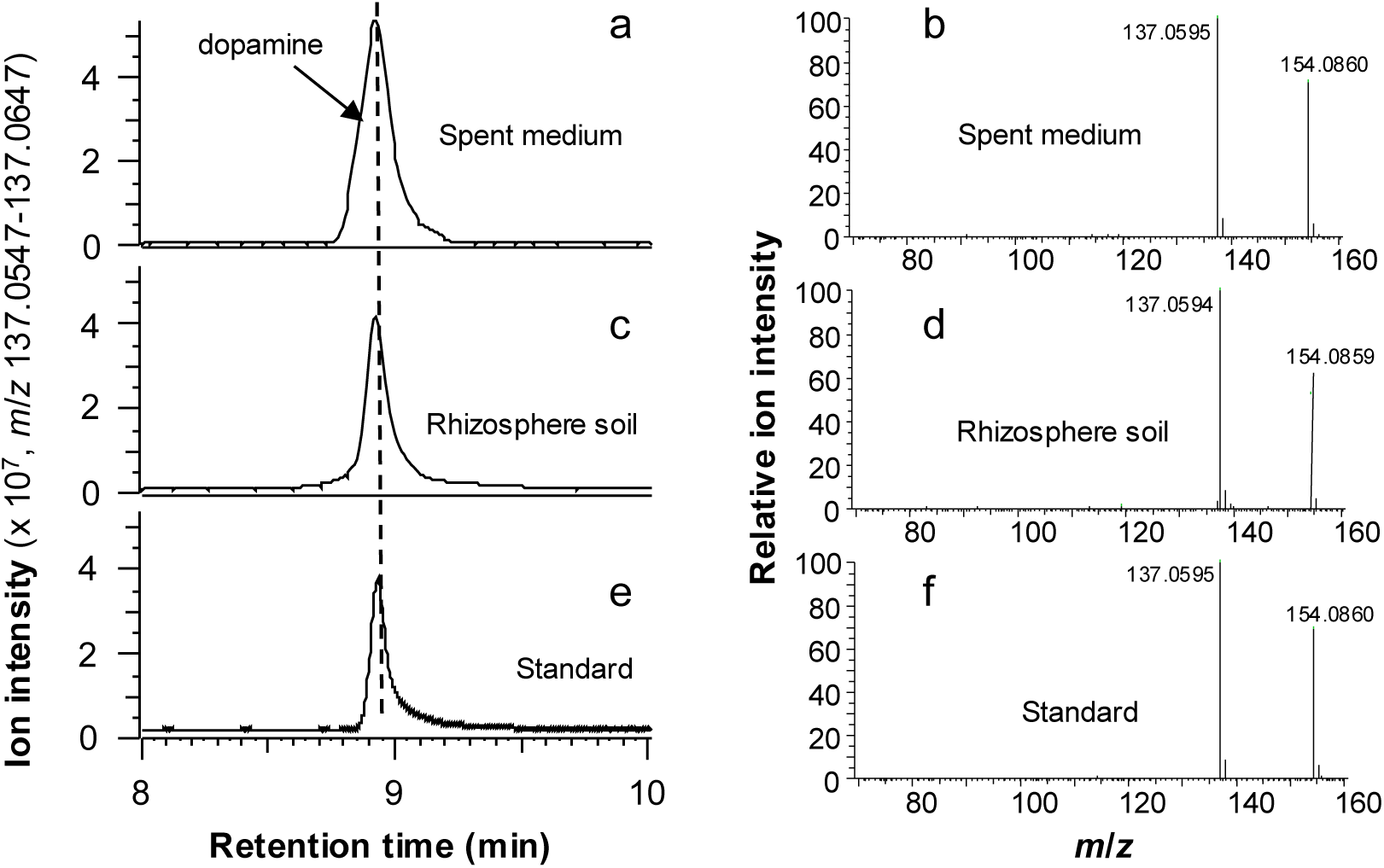
Identification of dopamine in *B. distachyon* root exudates and rhizosphere soil. Bd21-3 plants were grown in a hydroponic system and in soil. The growth medium and the rhizosphere soil were collected from 4-week-old plants for LC-MS/MS analysis. The left panel shows LC-MS/MS chromatograms of dopamine in the growth medium (**a**) and the rhizosphere soil (**c**) as compared to the standard (**e**). The right panel shows MS/MS fragmentation spectra of dopamine in the growth medium (**b**), the rhizosphere (**d**), and from the standard (**f**). Chromatograms and spectra are representatives of three biological replicates.

**Supplementary Fig. 9.**
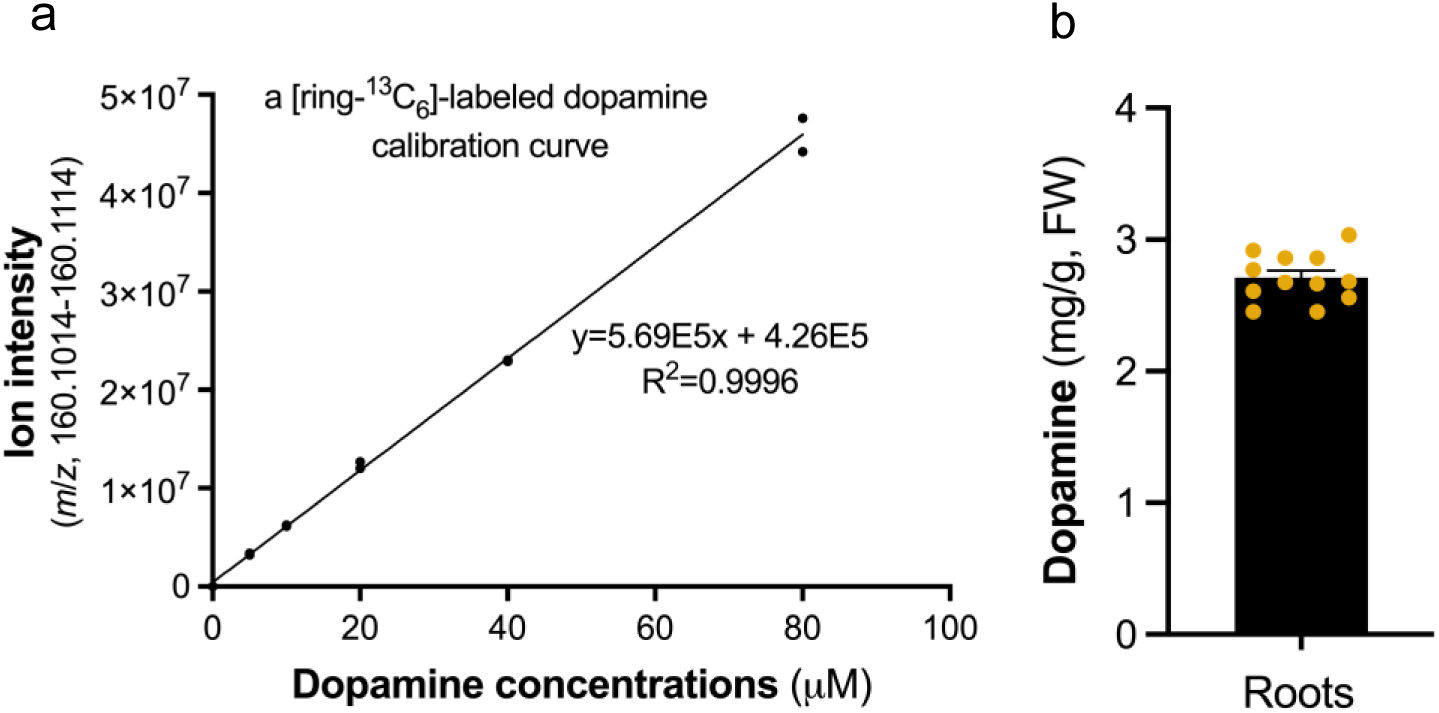
Quantification of dopamine in *B. distachyon* roots. **a** showing the internal dopamine calibration curve, which was derived based on the ion intensity of a [ring-^13^C_6_]-labeled dopamine (*m*/*z,* 160.1014-160.1114) over its concentration (*n* = 2). **b** showing endogenous dopamine levels (*m*/*z* 154.0812-154.0913) in *B. distachyon* roots measured based on the internal calibration curve (**a**) of a [ring-^13^C_6_]-labeled dopamine (Cambridge Isotope Laboratories). FW, fresh weight. Error bars represent mean ± SEM (*n* = 12).

**Supplementary Fig.10.**
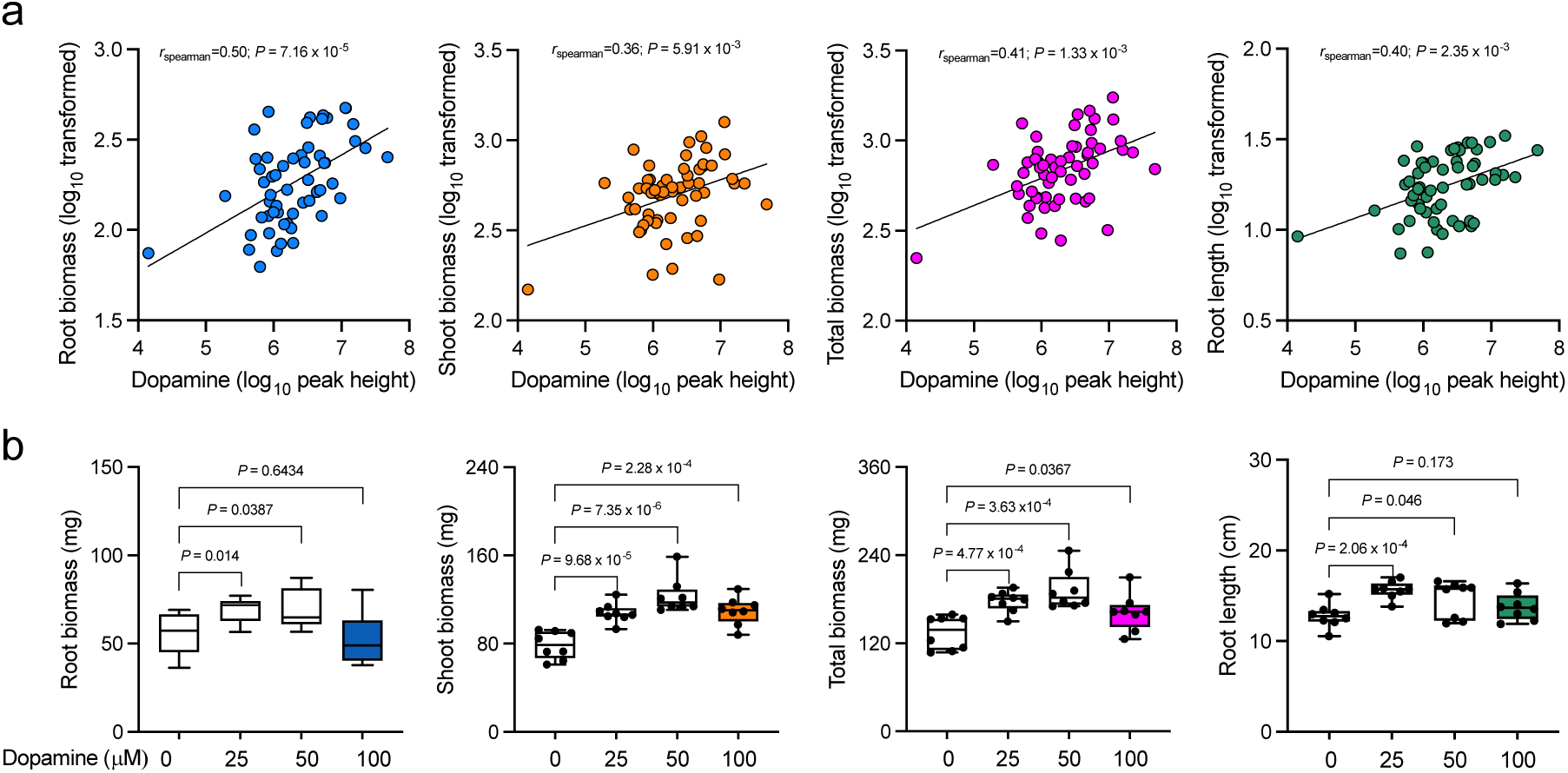
Dopamine has a positive effect on plant growth. **a** Spearman correlation between exudate dopamine levels and plant growth phenotypes, including root biomass, shoot biomass, total biomass, and root length. Log_10_ transformed data was used for correlation analysis (*n* = 55). **b** Dopamine could promote *B. distachyon* growth. *B. distachyon* Bd21-3 plants were grown in a hydroponic system supplemented with various concentrations of dopamine. Error bars represent mean ± SEM (*n* = 8). The *P-*value was calculated using the student’s *t-*test.

**Supplementary Fig. 11.**
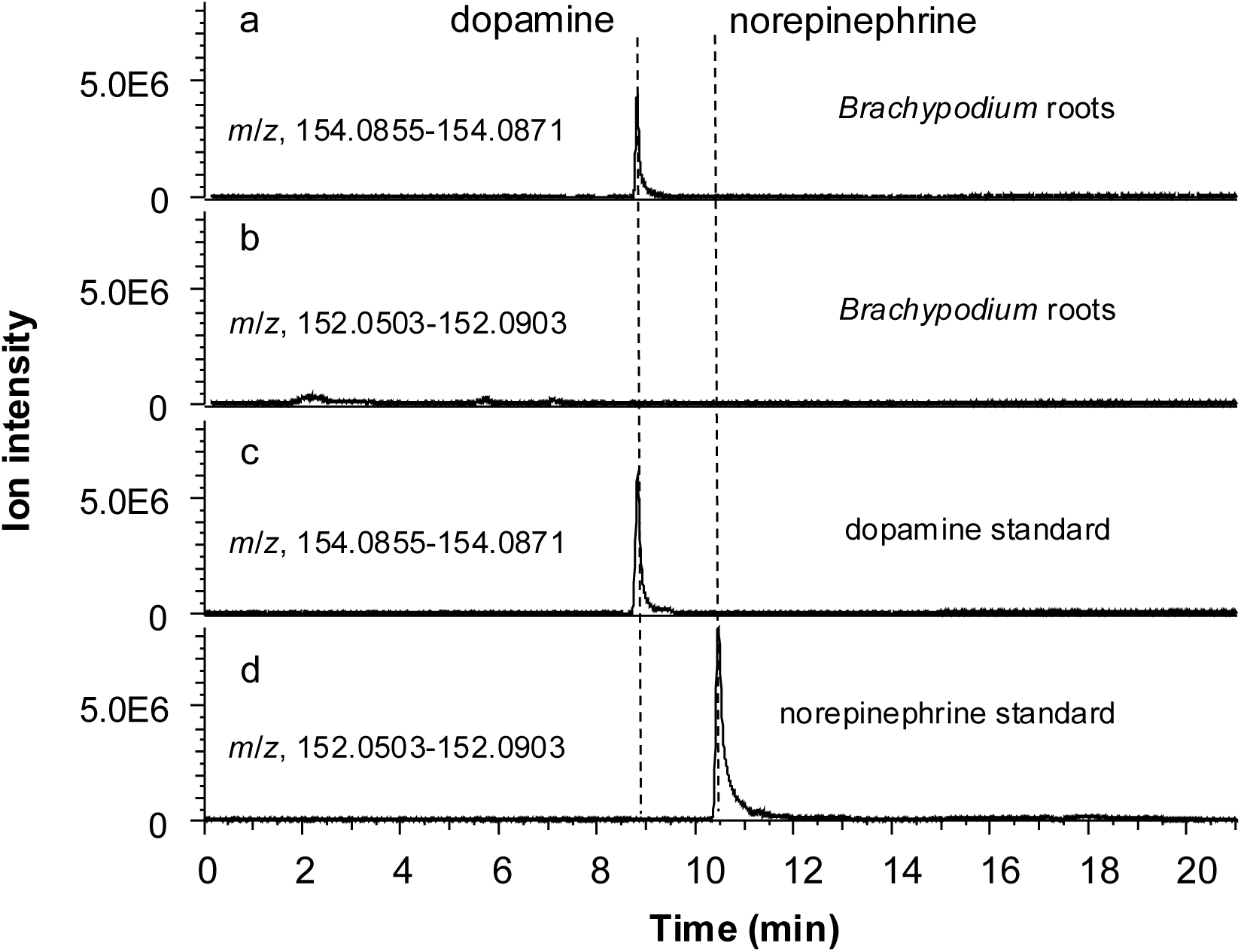
Identification of norepinephrine in *B. distachyon* roots. Bd21-3 plants were grown in a hydroponic system. The 4-week-old roots were used for LC-MS/MS analysis. The figure shows LC-MS/MS chromatograms of dopamine (**a** and **c**) and norepinephrine (**b** and **d**) in *B. distachyon* roots as well as standards. Chromatograms are representatives of three biological replicates.

**Supplementary Fig. 12.**
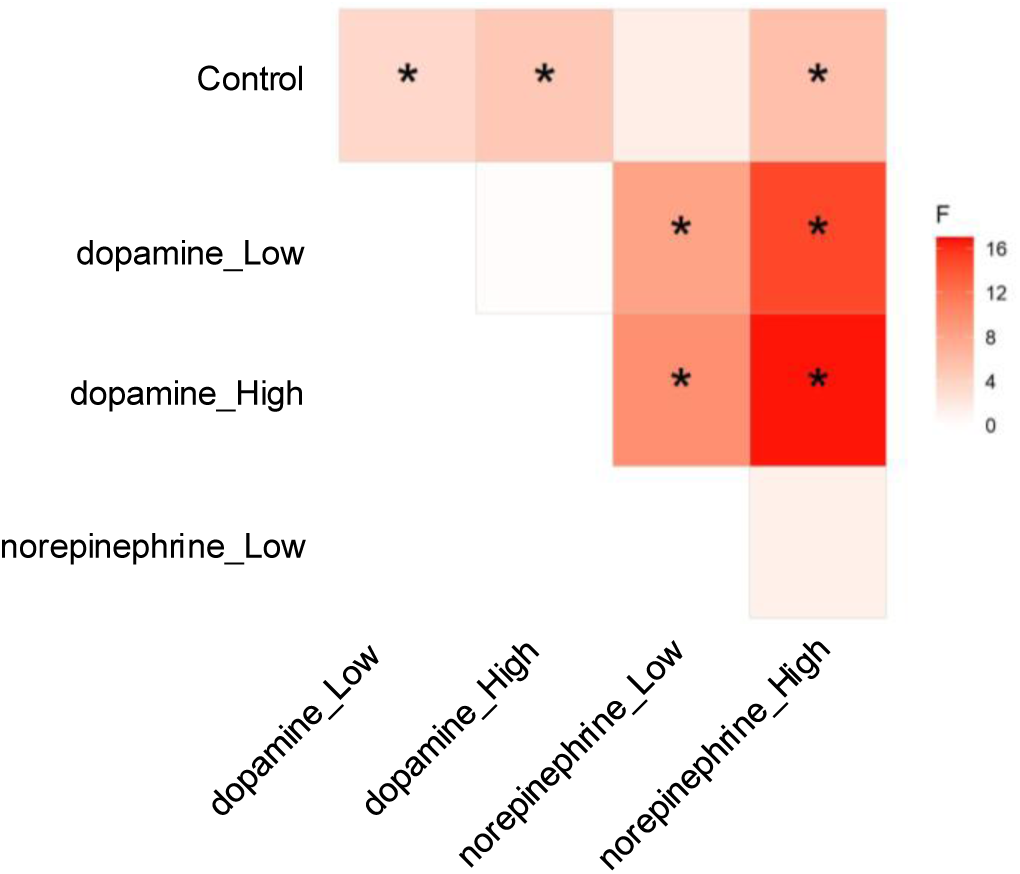
β-diversity of soil microbial communities in response to dopamine and norepinephrine. PERMANOVA was performed on the Bray-Curtis Dissimilarity among soil microbial communities in response to dopamine and norepinephrine treatments. The color of the matrix indicates the F score of the PERMANOVA between two treatments. Asterisks indicate significant β-diversity (*P*-value < 0.05).

**Supplementary Fig. 13.**
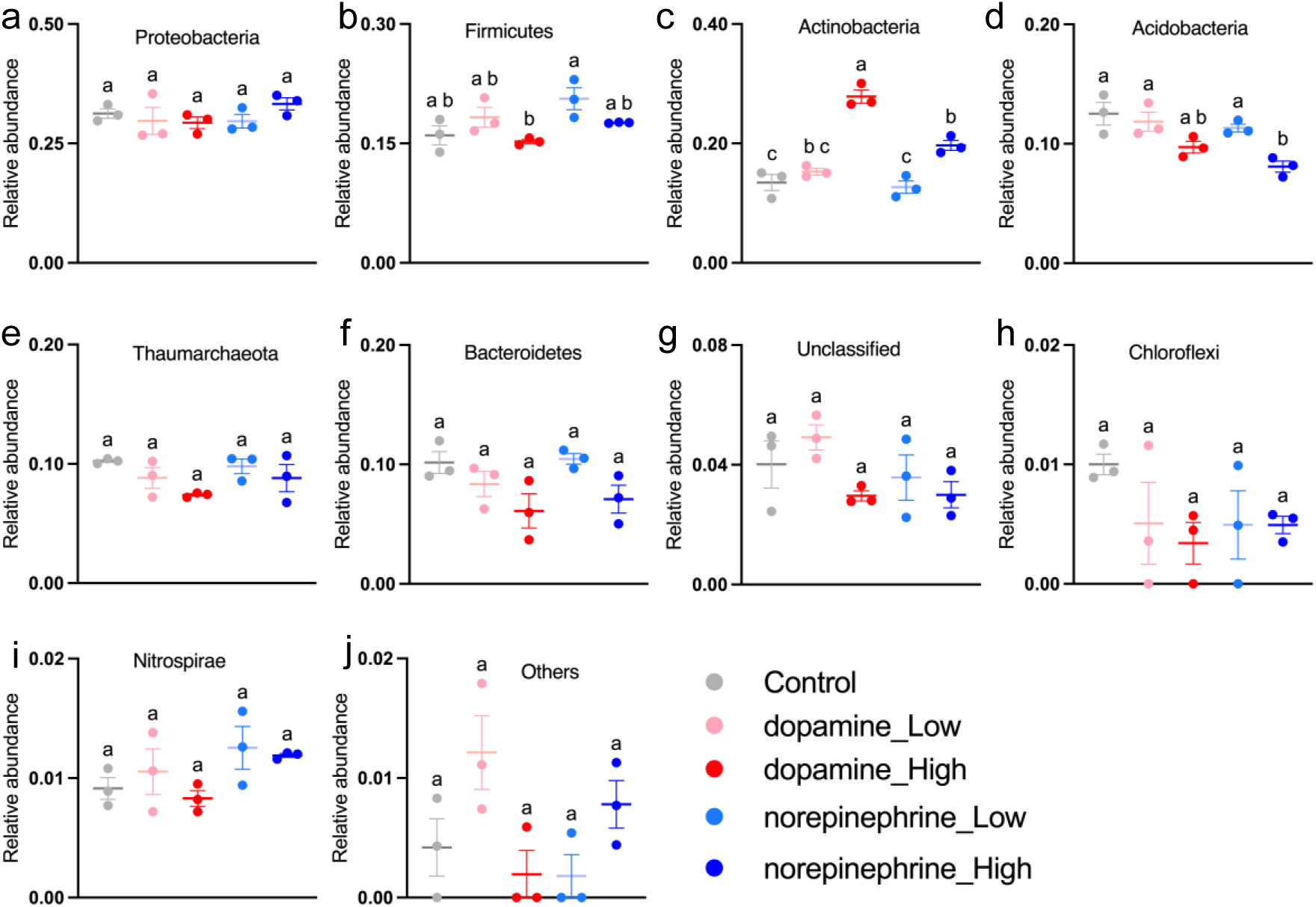
Dopamine enriched *Actinobacteria* in the soil at the phylum level. **a-j** showing bacterial phylum affected by dopamine and norepinephrine treatments. Error bars represent mean ± SEM (*n*=3). Within plots, different letters represent significant differences (one-way ANOVA followed by Tukey’s test corrections for multiple comparisons; *P* < 0.05).

**Supplementary Fig. 14.**
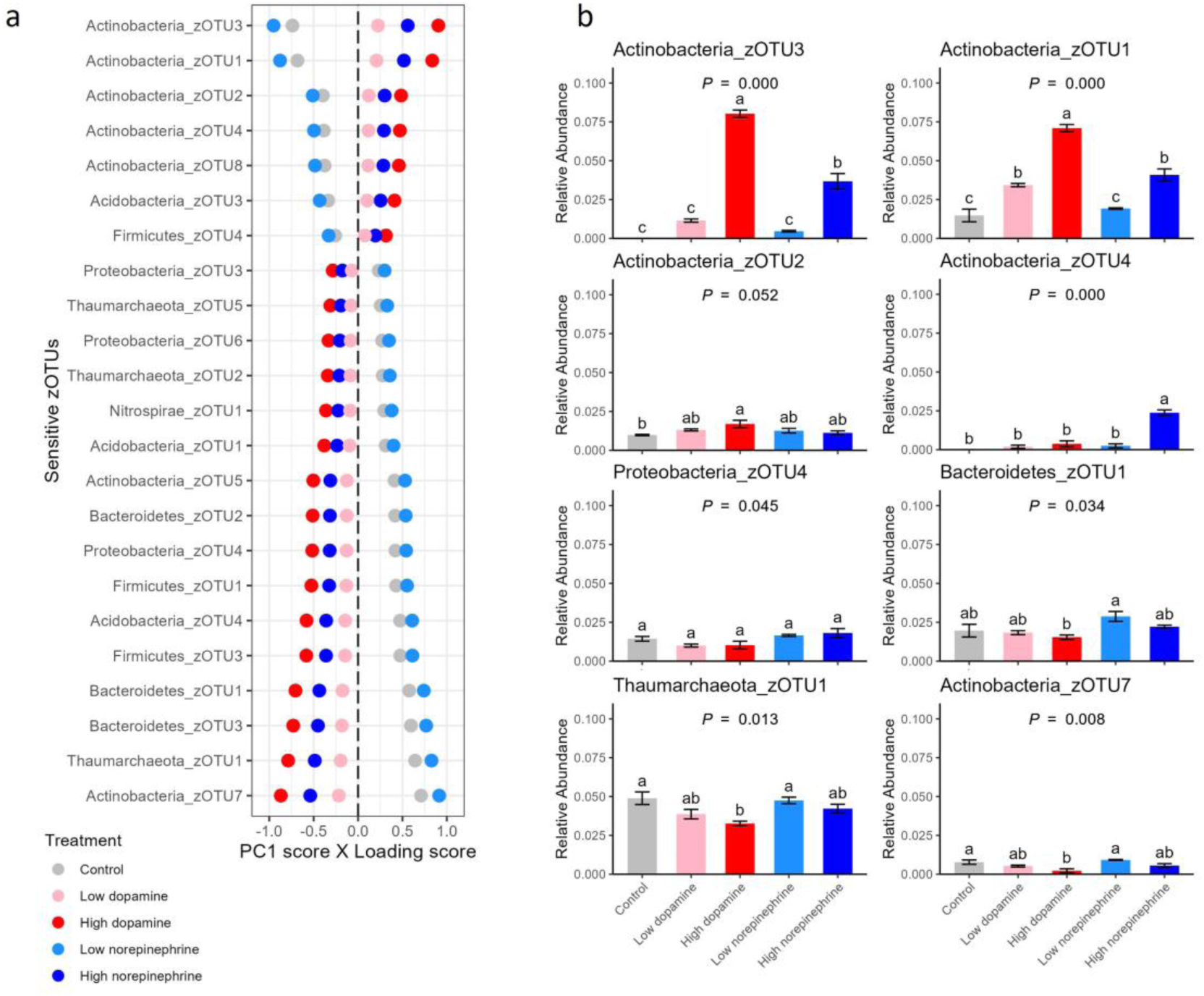
General trend of soil microbial response to dopamine and norepinephrine treatments. **a.** The relative responses of sensitive zOTUs to treatment are represented as their loading scores to principal component 1 (PC1) multiplied by the lsmeans of PC1 by treatment level. Analysis was done on zOTUs with relative abundance > 0.1% and those with loadings score to PC1 > 0.1 are displayed. b. The lsmeans of relative abundances of zOTUs with significant (*P*-value < 0.05; except Actinobacteria_zOTU2) ANOVA results by treatment levels. Error bars indicate the standard error of the mean (*n* =3), and lsmeans with different lower letters within each zOTU are statistically different (α = 0.05) based on Tukey’s post-hoc test.

